# Supporting-like cells constitute an alternative steroidogenic lineage conserved in amniotes

**DOI:** 10.64898/2026.05.25.726204

**Authors:** Iván Barberá-Aura, Wai-Yee Chung, Emilie Dujardin, Alicia Hurtado, Mathieu Galmiche, Chloé Mayère, Maëva Guy, Béatrice Mandon-Pépin, Vedran Franke, Yan Jaszczyszyn, Rafael D. Acemel, Serge Rudaz, Jennifer Mckey, Serge Nef, Eric Pailhoux, Darío G. Lupiáñez

## Abstract

In mammals, the production of sex hormones is widely considered to depend on the interstitial lineage of the gonad, which differentiates into Leydig cells in males or theca cells in females. However, certain mammalian species evidence gonadal steroidogenic activity prior to the specialization of these interstitial lineages, suggesting that alternative cell types may assume this function. Here we reveal a previously unrecognized role for supporting-like cells (SLCs), which can act as a major steroidogenic lineage during mammalian embryonic development. Through comparative single-cell transcriptomics, steroidomics and *in toto* organ imaging we find that in rabbits, SLCs not only contribute to the formation of gonadal rete structures, as described for other mammals, but also differentiate into a steroid-producing population. The steroidogenic program of SLCs is initially activated in both sexes but selectively maintained in ovaries, whereas in testes it is progressively replaced by that of interstitially derived Leydig cells. Evolutionary comparisons indicate that SLCs may represent an ancestral lineage that is homologous to the steroidogenic cells of non-mammalian species, which also derive from supporting precursors and share expression of cell fate regulators such as PAX2/8 and TBX1. Altogether, our findings redefine current models of gonadal lineages, revealing an unexpected plasticity in sex differentiation and exemplifying how distinct cell types can converge on analogous functions during evolution.

## INTRODUCTION

Sex hormones, which include progestogens, androgens and estrogens, are fundamental players for reproduction, physiology and behaviour. Sex hormones are mainly produced in the gonads, which contain specialized cell types that support steroidogenesis^1,2^. In mammals, the expression of the *Sex-determining region Y (SRY)* gene in XY embryos induces the specification of the supporting lineage into Sertoli cells^3,4,5,6^, which in turn promotes the differentiation of interstitial progenitors. This lineage will give rise to fetal Leydig cells, which produce androgens responsible for embryonic masculinization^7,8,9^. In XX embryos, which lack *SRY*, ovarian development is actively driven by ovarian-promoting factors that include the Wilms tumor suppressor (WT1) −KTS isoform and components of the WNT signaling pathway^10,11,12,13,14,15^. These factors induce the differentiation of the supporting lineage into granulosa cells, a key component of the ovarian follicles. Postnatally, granulosa cells promote the differentiation of the ovarian interstitial lineage into theca cells, the primary steroidogenic lineage, which play essential roles in follicle formation and function^16^. Thus, regardless of the sex, interstitially-derived cells are considered an indispensable component for sex hormone production, as they ensure the presence of the full enzymatic complement required for progestogen and androgen biosynthesis (STAR, CYP11A1, CYP17A1, HSD3B). In the ovary, granulosa cells cooperate with theca cells, but only for the final conversion of androgens to estrogens through the action of aromatase. Collectively, sex hormones are essential to maintain gonadal function and gametogenesis, while also exerting effects at the organismal level, including the development of secondary sexual characteristics and the modulation of behavior^17,18^.

Beyond these roles in reproduction and organismal fitness, sex hormones and in particular estrogens, can also play a central role in vertebrate sex determination. In fishes, estrogens are essential for the ovarian fate by antagonizing male pathway genes, with mutations affecting estrogen synthesis or receptor signaling often leading to female-to-male sex reversal^19,20^. In reptiles with temperature-dependent sex determination (TSD), such as turtles and alligators, estrogens can induce female development even at male-producing temperatures by activating estrogen receptors, particularly ESR1^21,22,23^. Consequently, the inhibition of aromatase, the enzyme that converts testosterone into estradiol, typically results in male development in amphibians, reptiles, and birds^24,25,26^. Altogether, these observations establish estrogens as conserved regulators of sex determination across major vertebrate lineages.

However, mammals represent a striking exception for this paradigm^27^. In this clade, sex determination is genetically controlled, with hormones mainly acting at later stages of differentiation. Intriguingly, estrogenic activity has been reported during the sex determination period for certain mammalian species. Early estradiol production or expression of *CYP19A1*, the gene encoding aromatase, has been described in fetal gonads from rabbits^28,29,30,31^, sheep^32,33^, goats^34^, cattle^35,36,37^, or humans^38^. Although loss-of-function mutant rabbits for *CYP19A1* are able to initiate female development, they show reduced ovarian size and aberrant cell proliferation^27^, resulting in a huge decrease of the ovarian reserve at puberty. These findings suggest that although estrogens are not primary determinants of sex in mammals, early steroidogenic activity may contribute to key aspects of gonadal development. Notably, estrogen activity occurs prior to the presence of theca cells^28,30,31^, the main steroidogenic cell type of the ovary, which only differentiates at postnatal stages after follicles are formed. This temporal gap suggests the existence of alternative cell types that may transiently assume the steroidogenic function in the early gonad. However, the molecular origin and evolutionary significance of this steroidogenic activity in mammals remain unexplored.

Advances in single-cell omics technologies have provided novel means to elucidate the cellular composition of developing gonads, redefining our understanding of the origin and relationships between cell lineages. A comprehensive single-cell atlas of the developing mouse gonad recently identified a lineage termed supporting-like cells (SLCs)^39^. These cells were shown to contribute to the formation of rete structures, which are epithelial networks connecting ovaries or testes with the mesonephros. In addition, SLCs contribute to a minor fraction of the gonadal pool of supporting cells^39^. Importantly, the SLC lineage appears to be evolutionary conserved across mammals, as it is also present in other species^40^. Beyond gonadal lineages, comparative single-cell analyses across vertebrate lineages have also suggested that cell functions may be more flexible than previously appreciated^41^. While the role of supporting cells in orchestrating sex determination and of the interstitial lineage on steroidogenesis are well accepted, studies in non-mammalian species have provided a more nuanced view. In species like chicken and turtle, prenatal steroidogenesis is largely driven by supporting cells, instead of the interstitial lineage^41,42^. Yet to what extent these alternative steroidogenic activities reflect ancestral mechanisms or species-specific features remains unclarified.

Here we investigate the molecular underpinnings of early mammalian steroidogenesis by studying the rabbit, a species in which this process takes place prior to the specialization of interstitially-derived lineages into steroid-producing cells^27^. By combining single-cell transcriptomics, steroidomics and *in toto* gonadal imaging analyses, we identify a previously unrecognized role for SLCs in steroidogenesis. At early developmental stages, SLCs acquire a sexually dimorphic identity and activate the steroidogenic program in both XX and XY gonads. In the testis, this steroidogenic program is progressively replaced by that of interstitial Leydig cells, whereas in the ovary SLCs remain as a major steroidogenic population during the entire prenatal development. Lineage reconstruction analysis revealed that SLCs follow two distinct cellular trajectories, one contributing to rete structures, as also described in other mammals^39^, and another originating steroidogenic cells. Comparative analyses with non-mammalian vertebrates reveal striking similarities with the supporting lineage of turtles, which also displays steroidogenic activity and shares the expression of lineage-specific markers like PAX paralogs or TBX1. Altogether, our findings reveal a latent steroidogenic potential within supporting gonadal lineages, highlighting the cellular plasticity underlying vertebrate sex development.

## RESULTS

### Supporting-like cells constitute a major cell population of the prenatal rabbit ovary

To elucidate the cellular composition of embryonic rabbit gonads, we performed single-cell RNA-seq experiments at 7 different developmental stages, generating a XX and XY gonadal atlas for the species *Oryctolagus cuniculus*. Our single-cell atlas includes gonadal undifferentiated (13 and 15 days *post-coïtum* (*dpc*); including adjacent mesonephros), sex-determining (17 and 19 *dpc*), and differentiated stages (21 *dpc*) from both sexes. Two additional stages of female gonads were included to study late embryogenesis (24 and 28 *dpc*) (Fig 1a). Altogether, we sequenced 195,607 cells that passed quality control filters. We integrated the data using dimensionality reduction and identified a total of 37 clusters (Supp Fig. 1a). As expected, cells from undifferentiated gonads of both sexes colocalized in the reduced embedding space, whereas those corresponding to subsequent XX and XY developmental stages occupied distinct spaces, consistent with the activation of sex-specific transcriptional programs (Fig 1b).

**Figure 1.**
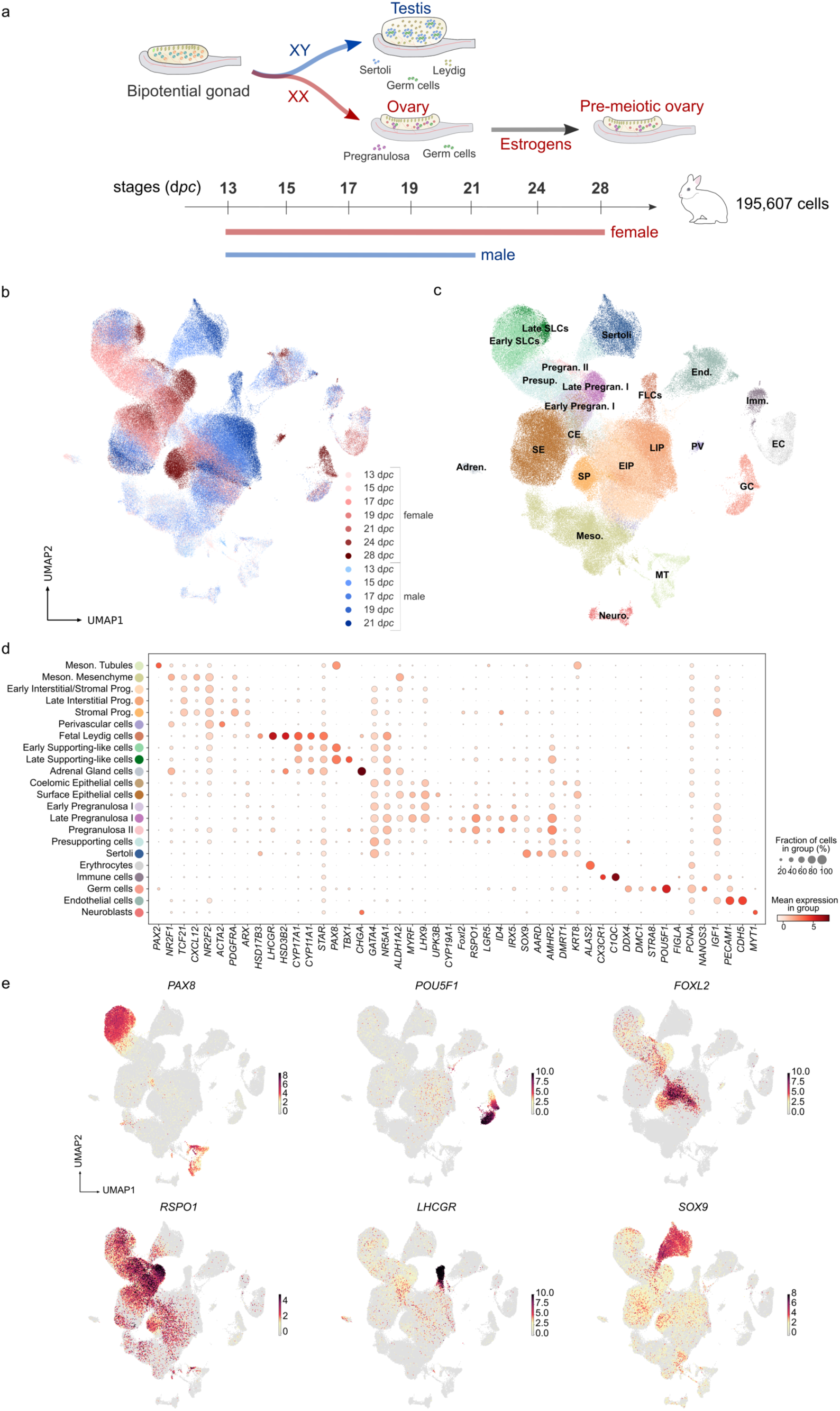
A single-cell atlas of rabbit gonadal development. a) Schematic representation of the gonad differentiation in rabbits and of the role of estrogens. Single-cell RNA-seq libraries were prepared for different stages encompassing the complete prenatal differentiation process. b) UMAP projection of the integrated samples coloured by sex and stages. c) UMAP projection coloured by cell types. d) Dotplot displaying the mean scaled expression of canonical gonadal and extragonadal markers per cell type. The dot size represents the fraction of cells in the cell type expressing the gene and the intensity of the color the mean scaled expression in the cell type. e) UMAP projection coloured by the scaled expression of the genes *PAX8*, *POU5F1*, *FOXL2*, *RSPO1*, *LHCGR* and *SOX9*. Early SLCs, early supporting-like cells; Late SLCs, late supporting-like cells; CE, coelomic epithelial cells; SE, surface epithelial cells; Presup., presupporting cells; Sertoli, Sertoli cells; Pregran. II, pregranulosa II cells; Early Pregran. I, early pregranulosa I cells; Late Pregran. I, late pregranulosa I cells; EIP, early interstitial/stromal progenitors; LIP, late interstitial progenitors; SP, stromal progenitors; FLC, fetal Leydig cells; PV, perivascular cells; Imm., immune cells; Meso., mesonephric mesenchymal cells; MT, mesonephric tubules; End., endothelial cells; GC, germ cells; EC, erythrocytes; Neuro., neuroblasts; Adren., adrenal gland cells

To assign identities to each cell cluster, we classified them according to known markers^39^. This initial annotation was then refined for the female and male samples separately, so that underrepresented cell types in the full dataset were correctly annotated (Fig 1c). Somatic cells were identified by the expression of *GATA4,* and expected mammalian gonadal cell types were present, including female-specific pregranulosa cells (*RSPO1*+, *LGR5*+), male-specific Sertoli (*SOX9*+) and fetal Leydig cells (*LHCGR*+)^43,44,45,46,47^. Supporting-like cells (SLC; *PAX8*+), a cell type recently identified in the mouse and human, were also present in both sexes^39,40^ (Fig 1b-e). As described for other species, we identified two distinct subpopulations of pregranulosa cells: Late Pregranulosa I, which closely resembles the Early Pregranulosa I cells found in earlier stages, and Pregranulosa II, which is only found in late stages 24 and 28 and characterized by prominent expression of the *FOXL2* gene^48^ (Fig 1d and e). Additionally, *FOXL2* expression was also observed in stromal progenitors. Interestingly, *HSD17B3* expression, which is required for converting androstenedione to testosterone in the testes, was not restricted to Sertoli cells as previously demonstrated in the mouse and human^49,50^, but it was also detected in fetal Leydig cells (Fig 1d). *POU5F1*, known for its role in maintaining pluripotency^51^, was broadly expressed in germ cells from both sexes in all stages. In contrast, *STRA8,* a marker regulating the onset of meiosis in response to retinoic acid signaling^52^, was exclusively detected in female germ cells from the latest stages (Supp Fig. 1b). This is consistent with the delayed start of meiosis in rabbit ovaries^53^. Mesonephric mesenchymal cells were identified by the expression of *NR2F1*, and the absence of *GATA4*. Mesonephric tubule cells featured the expression of *PAX8* and *PAX2*. Extragonadal cell types were also identified, including neuroblasts (*MYT1*+, *CHGA*+), immune cells (*C1QC*+, *CX3CR1*+), adrenal gland cells (*CHGA*+, *HSD3B2*+) and erythrocytes (*ALAS2*+) among others (Fig 1d).

To gain further insights on sex-specific differences in gonadal composition, we extracted cell proportion for each stage and cell cluster (Fig 2a). This analysis revealed a higher proportion of the interstitial lineage in the late male stages in comparison with their female stromal counterpart. More specifically, interstitial progenitors and fetal Leydig cells, which account for more than half of the cells of the developing testes at stages 17, 19 and 21 *dpc*. This is consistent with higher proliferative state of the male gonad, associated with the activation of the androgenic pathway during testicular development in mammals^54,55,56,9^. Notably, we observed a higher proportion of SLCs in the female somatic gonad, compared to the male counterpart, particularly during undifferentiated and sex-determining stages (15-19 *dpc*; Fig 2a). To test for an enrichment of SLCs between sexes, we performed cell-neighborhood differential abundance analysis. This analysis generates small subpopulations of cells on the k-nearest neighbor (KNN) graph and tests for differential abundance between experimental conditions across those subpopulations. The analysis revealed 392 significantly enriched (SpatialFDR < 0.1) SLCs neighborhoods, that is small groups of cells belonging to that cell type, in rabbit female cells against only 17 neighborhoods significantly enriched in male cells (Fig 2b). A differential abundance analysis on mouse SLCs, which were described as a reduced population in the gonad^39^ (Fig 2c), confirmed no apparent sex bias (Fig 2d), with only 6 significantly enriched SLC neighborhoods in female samples. Similar results were observed for human SLCs at 8.6 post-conception week (PCW) (approximately equivalent to mouse 13.5 *dpc*) with the SLCs being a minority population with a slightly male bias (Supp Fig. 2 a-b). To evaluate whether the high proportion of SLCs in the rabbit was due to a high proliferative state of these cells, cell cycle scores were computed for each cell based on the expression of cell cycle markers. These analyses suggest that the vast majority of the SLCs remain quiescent in G1 independently of the sex and stage (Supp Fig. 1c) similarly to what was described for the mouse SLCs^39^. Even though the proportion of SLCs is greater in the female than in the male, no remarkable differences regarding the cell cycle state were observed between the female and male SLCs (Supp Fig. 1d).

**Figure 2.**
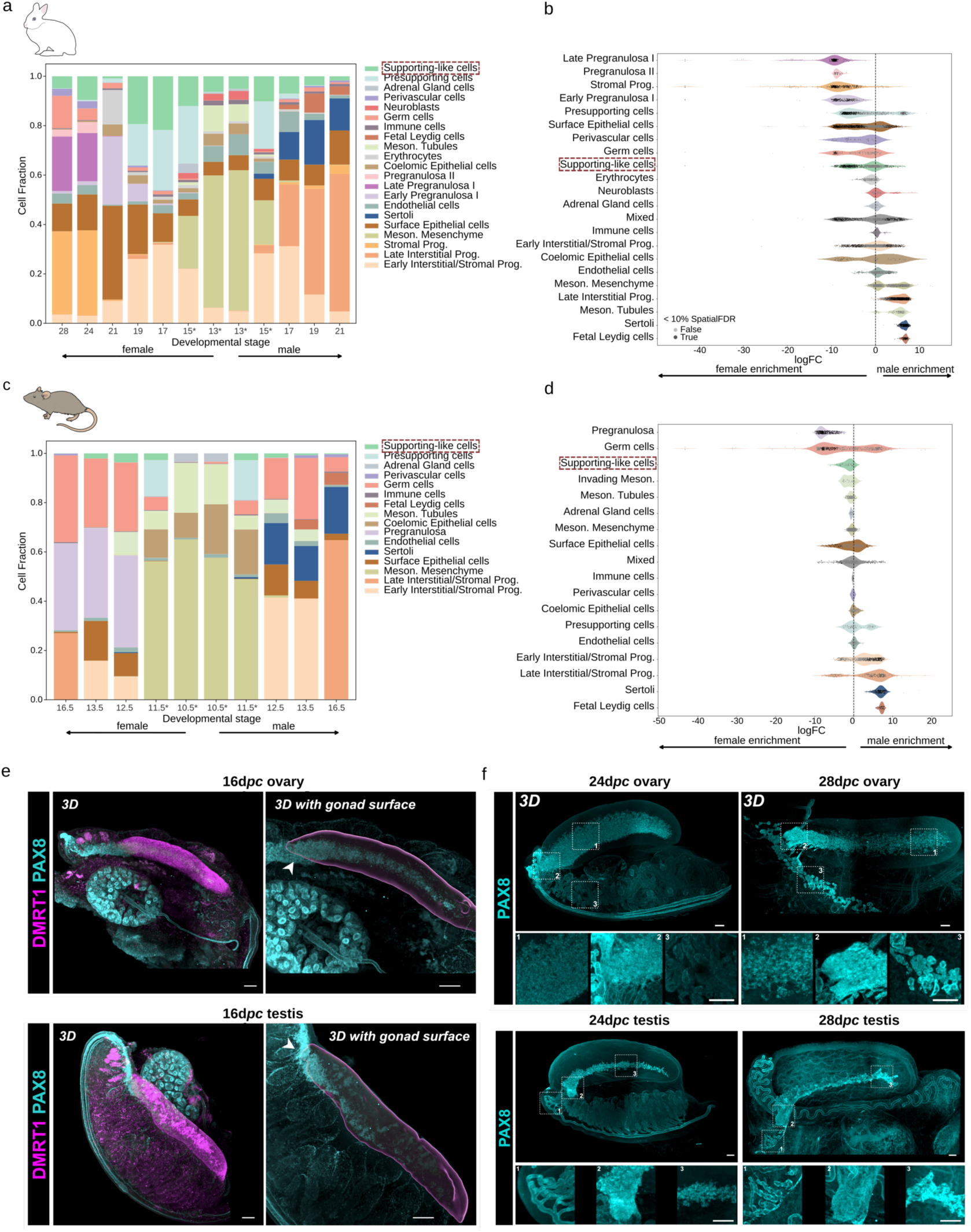
SLCs display differences in abundance across sexes and species. a) Barplot showing cell type proportions per sex and stage in rabbits. Adjacent mesonephros tissue was included for 13 and 15 *dpc* female and male samples (asterisk). b) Violin plot showing the logFC of the milo neighbourhoods per cell type of rabbit gonads. A negative logFC means that the neighbourhood is enriched in female cells and a positive logFC in male cells. Neighbourhoods displaying significant differential abundance between sexes (SpatialFDR < 0.1) are shown in dark grey, whereas not significant ones are represented in light grey. c) Barplot showing cell type proportions per sex and stage in mouse. d) Violin plot showing the logFC of the milo neighbourhoods per cell type of mouse gonads. e) *In toto* immunofluorescence images of ECi-cleared prenatal rabbit urogenital complexes at 16 *dpc*. Top row, ovaries and bottom row, testes. Left panels show the native immunofluorescence signal for DMRT1 (magenta) and PAX8 (cyan)s. Right panels show staining for PAX8 and 3D segmentation of the DMRT1+ domain, which corresponds to the gonadal surface. f) *In toto* immunofluorescence images of PAX8 (cyan) expression in ECi-cleared prenatal rabbit urogenital complexes at 24 *dpc* (left) and 28 d*pc* (right). Boxed areas correspond to magnified views of different anatomical positions of the urogenital complex, including gonad (box 1), gonad-mesonephros border (box 2) and mesonephros (box 3). Top row, ovaries and bottom row, testes. Scale bars, 100µm.

To examine the spatial distribution and dynamics of SLCs, we performed *in toto* imaging of ethyl-cinnamate (ECi)-cleared whole rabbit gonads and immunohistochemistry analyses on histological sections. PAX8+ SLCs were detected in early gonads (16 *dpc*) at the gonad–mesonephros interface in both sexes (Fig 2e), consistent with earlier findings^57^. Yet, this distribution differs notably between sexes as sex differentiation progresses. At 24 *dpc*, while SLCs remain at a similar position in the developing testis, they become a large compartment in the female gonad, occupying a considerable proportion of the medullary space (Fig 2f). Further analyses revealed that FOXL2+ cells were confined to the cortex and did not overlap with *PAX8* expression (Supp Fig. 3). Collectively, these analyses demonstrate that rabbit SLCs display a female-specific enrichment, particularly during sex-determining stages, a pattern that is distinct to what has been described in other mammalian species, such as mouse and human.

**Figure 3.**
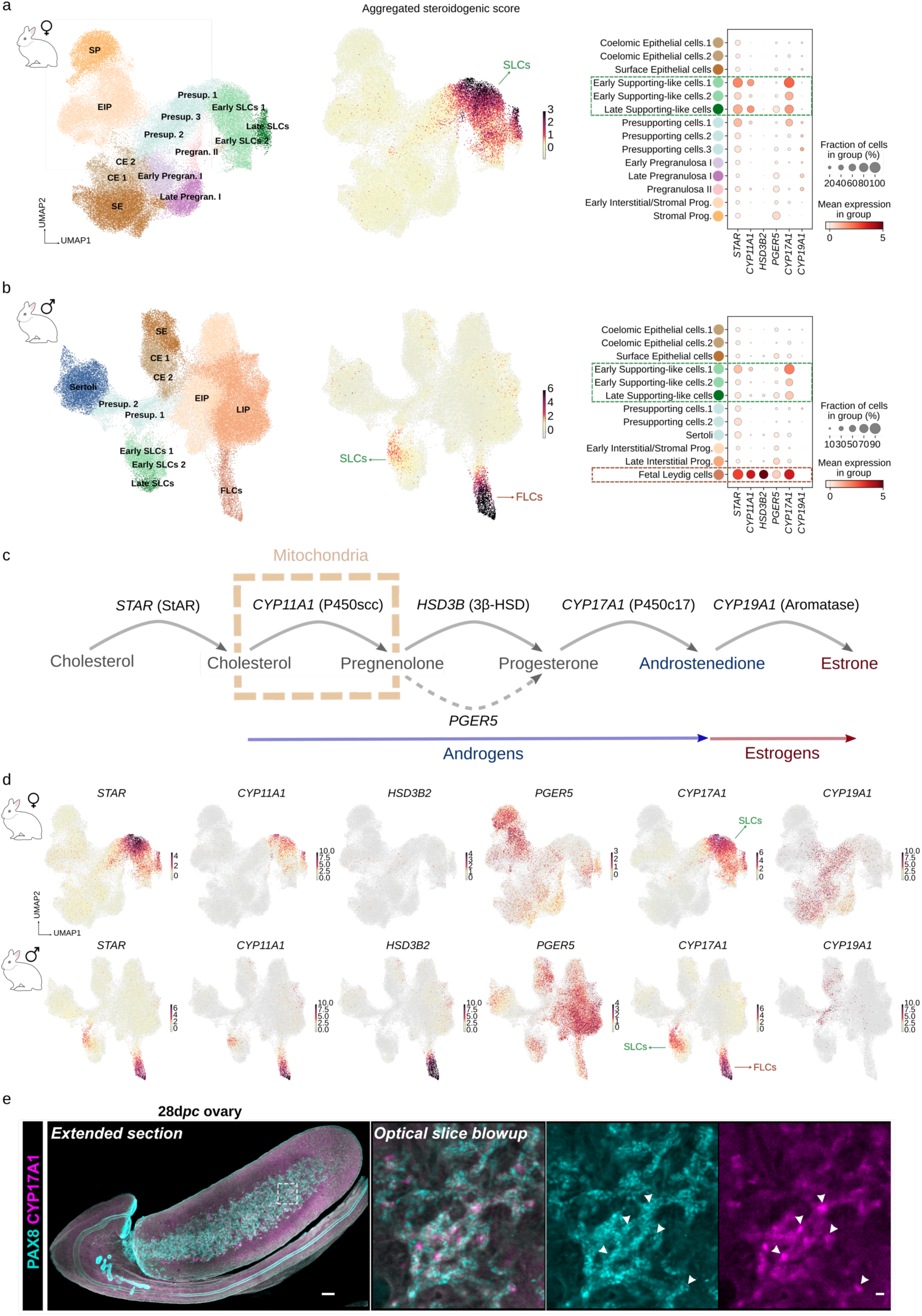
SLCs mediate steroidogenesis in the prenatal rabbit gonad. a) Left: UMAP projection of female somatic gonad coloured by cell type. Middle: UMAP coloured by the average expression of the steroidogenic genes *STAR*, *CYP11A1*, *HSD3B2* and *CYP17A1*. Right: Dotplot displaying the expression of the steroidogenic genes, in the cell types from the female somatic gonad. b) Left: UMAP projection of male somatic gonad coloured by cell type. Middle: UMAP coloured by the average expression of the steroidogenic genes *STAR*, *CYP11A1*, *HSD3B2* and *CYP17A1*. Right: Dotplot displaying the expression of the steroidogenic genes, in the cell types from the male somatic gonad. c) Schematics of sex hormone synthesis in the gonad. *PGER5* is indicated as candidate gene to compensate for the absence of *HSD3B2* expression in SLCs d) UMAP projections of female (upper panel) and male (lower panel) somatic gonads coloured by the scaled expression of the steroidogenic genes *STAR*, *CYP11A1*, *HSD3B2*, *PGER5*, *CYP17A1* and *CYP19A1*. Note that *PGER5*, although expressed in different cell types, also shows expression in SLCs e) *In toto* iImmunofluorescence image of ECi-cleared prenatal rabbit ovary at 28 *dpc*. Left panel shows a full extended section in the XY plane. Right panel corresponds to a single optical section magnified view of the boxed area. PAX8 is labelled in cyan, CYP17A1 in magenta. Scale bars: left, 100µm, right, 10µm. Early SLCs, early supporting-like cells; Late SLCs, late supporting-like cells; CE, coelomic epithelial cells; SE, surface epithelial cells; Presup., presupporting cells; Sertoli, Sertoli cells; Pregran. II, pregranulosa II cells; Early Pregran. I, early pregranulosa I cells; Late Pregran. I, late pregranulosa I cells; EIP, early interstitial/stromal progenitors; LIP, late interstitial progenitors; SP, stromal progenitors; FLC, fetal Leydig cells.

### Supporting-like cells mediate steroid synthesis during early rabbit gonadogenesis

Female rabbit gonads synthesize estrogens during early developmental stages, with estradiol reported to be present at 18 *dpc* and peaking around 20 *dpc*^28,30,31,27^. To identify the cell type responsible for steroid synthesis in the rabbit gonad, we examined the expression of key steroidogenic genes. We specifically focused on *STAR*, *CYP11A1*, *HSD3B2* and *CYP17A1*, genes encoding enzymes of the androgen-synthesis pathway, and on *CYP19A1*, which encodes aromatase, responsible of converting androgens into estrogens irreversibly. We subsetted the full dataset to focus on the somatic compartment and checked expression patterns in female and male cells separately (Fig 3a-b, Supp Fig. 4). Interestingly, all androgen-synthesis genes, except *HSD3B2*, were expressed in SLCs of both female and male rabbit gonads (Fig 3a-d, Supp Fig. 4a-b). *In toto* imaging analyses further confirmed the colocalization between SLC and steroidogenic marker genes (Fig. 3e). The expression of these genes was diminished at later stages in male SLCs, while remaining highly expressed in female SLCs throughout embryonic development (Supp Fig. 4c). The progressive inactivation of steroid genes in male SLCs was concomitant with the differentiation of fetal Leydig cells around 17-19 *dpc*, which overtake androgen synthesis, as also described for other mammals (Supp Fig. 4c). Notably, the expression of *HSD3B2* was exclusively restricted to fetal Leydig cells and not observed in SLCs. We also observed that *CYP19A1* was not expressed in female SLCs, but broadly distributed throughout the conventional supporting cell lineages, as previously shown^27^. As such, the estrogen synthesis process in the female rabbit gonad seems to be coordinated by different cell types: the SLCs that synthesize androgens, and the supporting lineage converting those to estrogens. This coordinated pattern was also described for non-mammalian vertebrates that synthesize estrogens during early gonadal development, implying a degree of evolutionary conservation^58,41,42^.

**Figure 4.**
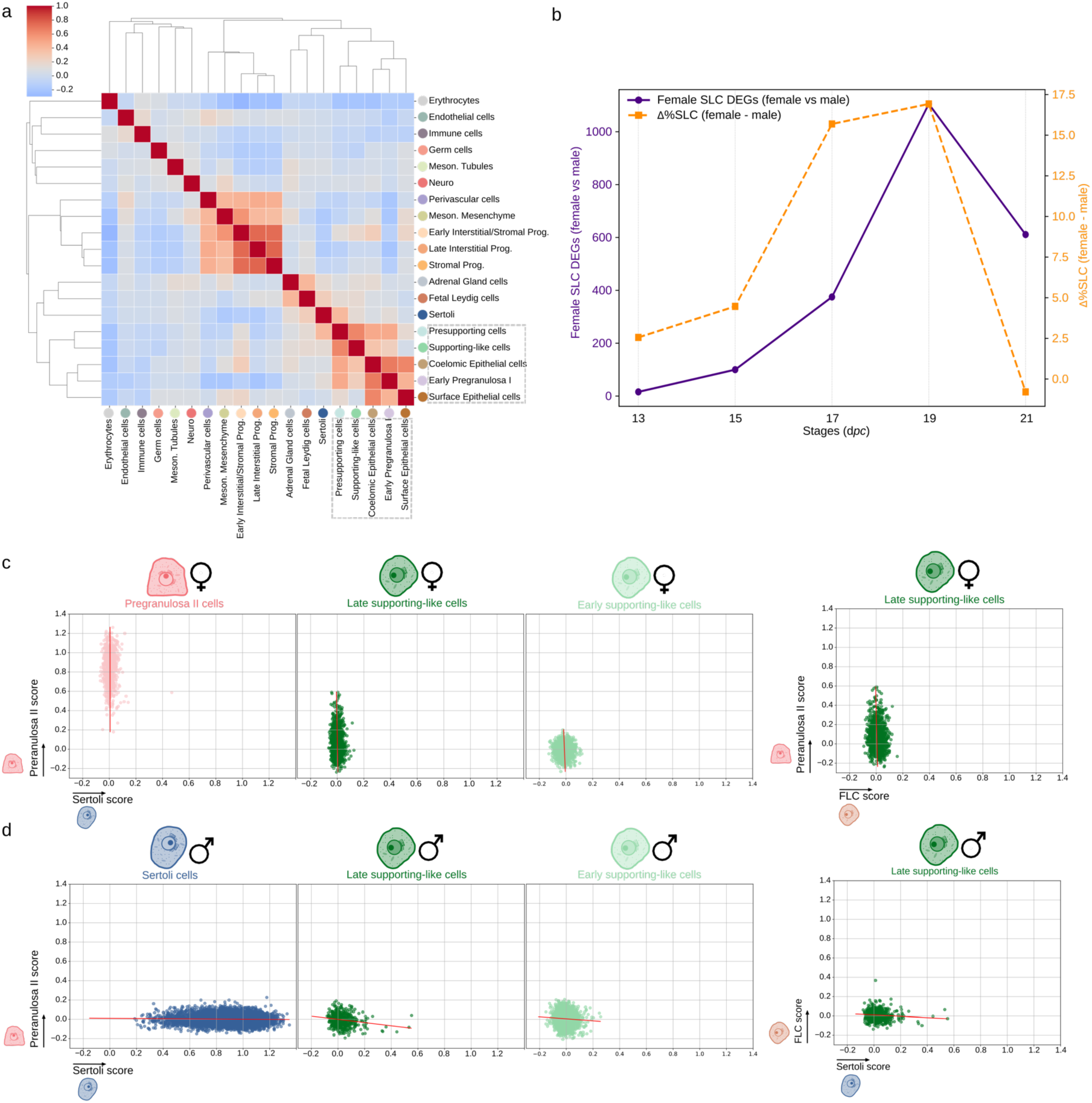
Transcriptional characterization of SLCs. a) Heatmap displaying the correlation between transcriptomes of all annotated cell types in rabbit gonads. Dashed square indicates the cell types showing more similarities with SLCs b) Lineplot highlighting the increase in the proportion of female/male SLCs per stage (orange), and the number of upregulated genes in the female SLCs vs male SLCs (purple). c-d) Scatterplots showing the pregranulosa II vs Sertoli scores (left plots), Pregranulosa II vs FLC scores (top right plot) and FLC vs Sertoli scores (bottom right plot) from LASSO models for female (c) and male (d) cell types.

Given the lack of *HSD3B2* expression in SLCs and its requirement for completing the androgen-synthesis pathway, we explored the steroid composition of the developing rabbit gonad. For this purpose, we performed extended steroid profiling through liquid chromatography – tandem mass spectrometry (LC-MS) according to a published protocol^59^ in bulk fetal rabbit gonads at 21 *dpc* and 28 *dpc*. A total of 173 steroid metabolites were targeted in the extended steroid profiling with relative quantification, including 9 major steroids with absolute quantification (Supp. Table 1). In male fetal gonads, 39 distinct steroid species were detected at 21 *dpc*, and 35 at 28 *dpc* (Supp Table 2). The male gonadal steroidome was dominated by androgens at 21 *dpc*, where they represented 18 of the 39 identified steroids in testes. Testosterone was particularly abundant (411 ± 42 ng/g tissue; Supp. Table 3). All testicular androgen concentrations were markedly lower at 28 *dpc* compared to 21 *dpc* (from 16- to 52-fold, Supp Fig. 5a). In female fetal gonads, a lower number of steroid metabolites could be detected (e.g., 13 at 21 *dpc*, and 23 at 28 *dpc*; Supp Table 2). Androgens were absent from female gonads, with the noteworthy exception of dehydroepiandrosterone (DHEA), the precursor of androgen synthesis, which is a CYP17 product. Yet most of its downstream products in the androgen synthesis pathway, usually requiring HSD3B, could not be identified in female fetal gonads. At 28 *dpc*, the androgen 11-ketotestosterone (11-KT) was identified in fetal ovaries, although none of its precursors could be detected, suggesting a non-gonadal origin. In both fetal testes and ovaries at 21 and 28 *dpc*, estrogens (that are produced from androgen precursors) could not be detected. This is explained by the combination of their very low endogenous concentrations and the low detection sensitivity of LC-MS for this group of hormones, which is commonly acknowledged in the field^60,61^. At 21 *dpc*, 9 different progestogens were detected in fetal testes, versus 5 in fetal ovaries. Interestingly, 3 out of these 5 progestogens (Pregnenolone and its 17α- and 7α-hydroxylated metabolites) were precursors of the steroid biosynthesis pathway that do not require the expression of HSD3B. However, progesterone could also be detected, though at low concentrations (0.064 ± 0.009 ng/g tissue; Supp. Table 3). Progesterone levels were thus 1000-fold lower than pregnenolone levels (60 ± 8 ng/g, Supp. Table 3). While in line with the absence of HSD3B in fetal ovaries (Supp Fig. 5b), these results also suggest the existence of an unexplored source for these minimal amounts of progesterone in the female rabbit gonad, outside the canonical HSD3B-based pathway converting from pregnenolone. Therefore, we examined our datasets for other members of the HSD gene family that may compensate for the absence of *HSD3B2* expression in female ovaries. Yet, the other *HSD3B* gene annotated in rabbits, *HSD3B1,* was also not expressed in SLCs at any stage of female gonadal development (Supp Fig. 6a). By expanding our analysis to other steroid-related genes, we found a *AKR1C* family member displaying an expression pattern that overlapped with the steroidogenic temporal window (Supp Fig. 6b-c). In the rabbit, this gene was annotated as *PGER5* (prostaglandin-E(2) 9-reductase-like) but exhibited the highest protein sequence similarity (∼80%) with human HSD17B5, which is encoded by *AKR1C3* (Supp Fig. 6d). Interestingly, the aldo-keto reductase superfamily (AKR) of humans has been proven to display 3β-hydroxysteroid dehydrogenase activity^62^, which is typically performed by the HSD3B family. *PGER5* expression increases from 17 *dpc* onwards, being slightly higher at 19 *dpc* and then at 24 and 28 *dpc* (Fig3 a,b,d and Supp Fig. 6b-c), which coincides with the increase of expression of *CYP19A1*. Other *HSD* and *AKR1C* genes showed either no compatible temporal expression pattern with the synthesis of estrogens, or a function that did not resemble that of *HSD3B2*. These results suggest that pregnenolone to progesterone conversion, which is a critical step for steroid synthesis, takes place in the developing rabbit ovary. Yet, this conversion occurs with low efficiency, likely due to the activity of a non-canonical gene with hydroxysteroid dehydrogenase, *PGER5*.

**Figure 5.**
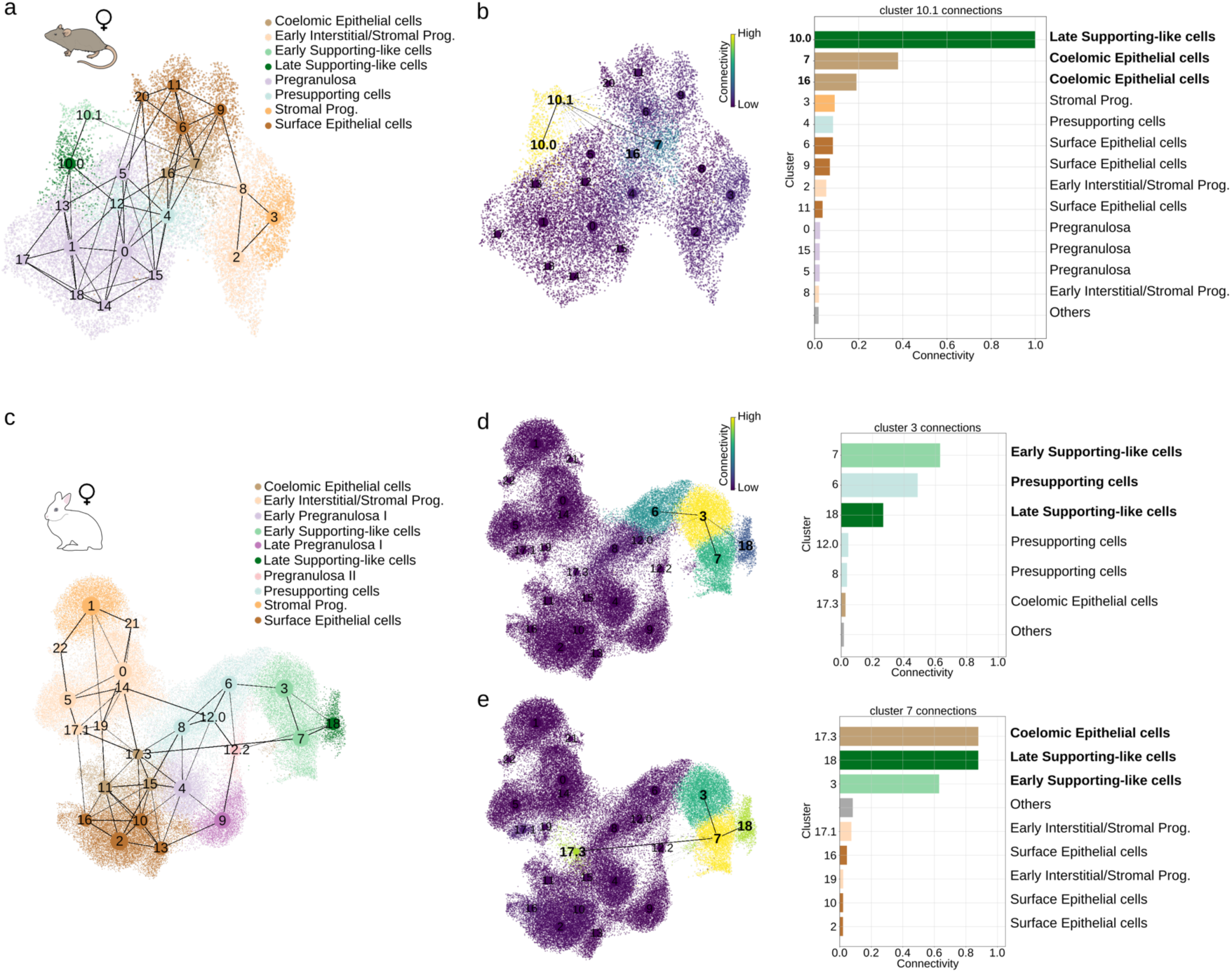
Developmental origin of rabbit SLCs. a) PAGA connectivity graph of female mouse somatic gonads. Additional subclustering was performed for some coelomic epithelium and SLCs clusters to ensure better resolution. The width of the connections represents the connectivity inferred by PAGA and potentially reflects cell type similarities. b) PAGA connections restricted to the early SLCs cluster (cluster 10.1). The intensity of the colour in the UMAP (left plot) represents the connectivity of the clusters to the cluster 10.1 and it is equivalent to the length of the bars in the barplot (right plot). c) PAGA connectivity graph of female rabbit somatic gonads. d-e) PAGA connections restricted to the subclusters of early SLCs. Cluster 3 (d) is strongly connected to presupporting cells and cluster 7 (e) to the coelomic epithelium.

**Figure 6.**
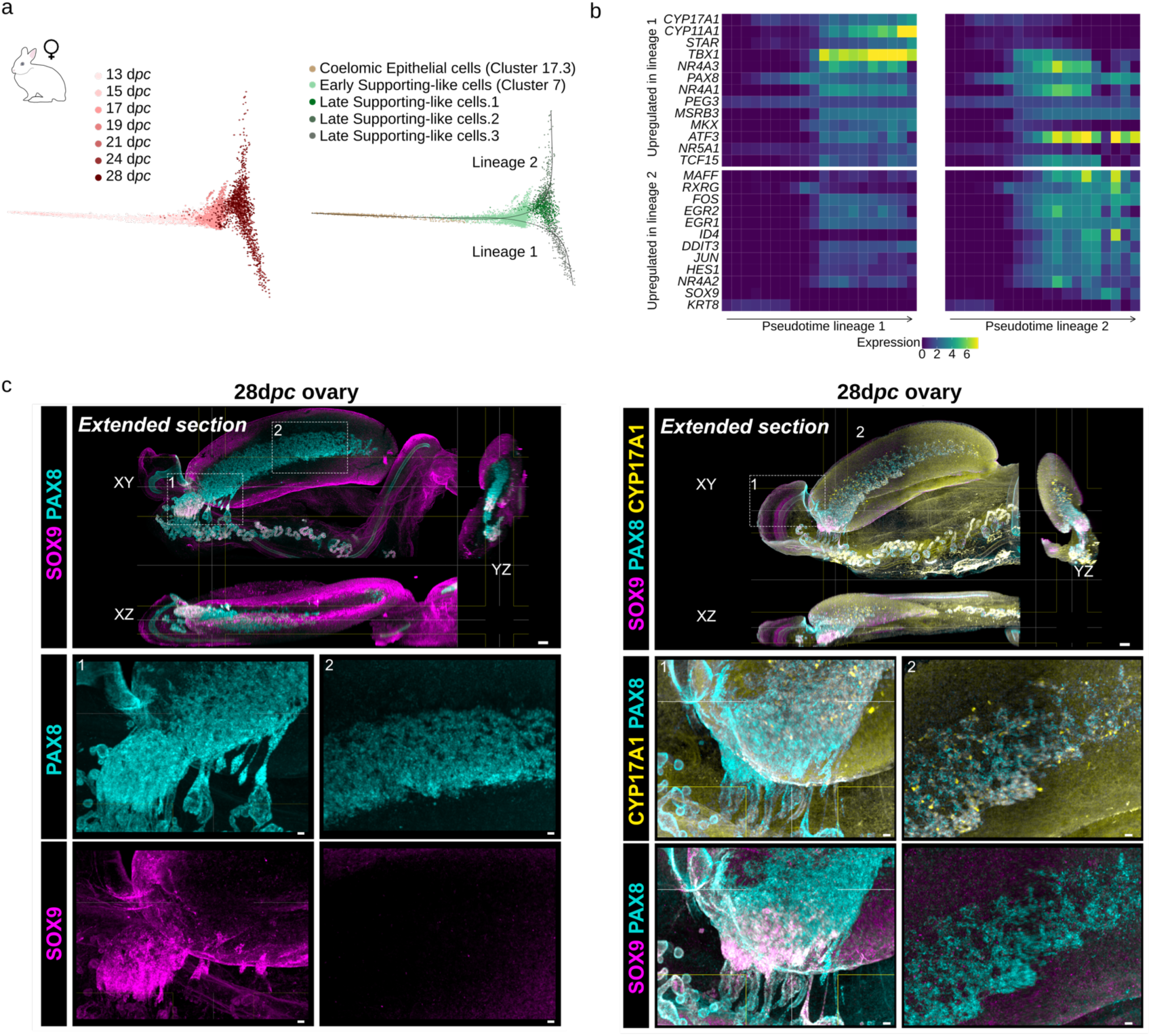
Rabbit SLCs differentiation trajectories and terminal states. a) Diffusion maps of PAGA highly connected clusters starting from the coelomic epithelium (cluster 17.3) and ending in late SLCs of the female rabbit. Left: diffusion map coloured by stages type. Right: diffusion map coloured by cell types. New subpopulations of late SLCs were identified. Slingshot was used to order the cells along two lineages and to compute a pseudotime. b) Heatmap showing gene expression trends along the slingshot lineages for the androgen-synthesis genes, upregulated genes in at least one of the lineages, *SOX9* and *KRT8*. Cells were separated in 20 blocks based on their pseudotime values. The colours represent the mean expression value in each block. c) *In toto* Immunofluorescence images of ECI-cleared prenatal rabbit ovaries at 28 *dpc*. PAX8 is labeled in cyan, SOX9 in magenta and CYP17A1 in yellow. Colocalization of PAX8+ and SOX9+ cells is observed near the border with the mesonephros (box 1), whereas colocalization of PAX8+ and CYP17A1+ cells is observed in the medullary region (box 2). Scale bars: top, 100µm; bottom, 20µm.

Altogether, our results demonstrate that estrogen synthesis in the rabbit early ovary is carried out primarily by SLCs, in cooperation with the supporting lineage for the final conversion step. In males, the steroidogenic program of SLCs is also triggered but progressively replaced by fetal Leydig cells, whereas in females it remains active throughout prenatal development.

### Supporting-like cells are transcriptionally distinct from other gonadal steroidogenic lineages

Given the steroidogenic nature of rabbit SLCs, we explored whether this cell type displayed a similar transcriptional profile to other steroid-producing lineages of the gonad. For this purpose, we calculated the mean scaled expression of each gene per cell type and used it for computing the Pearson correlations across the full dataset (Fig 4a). Overall, SLCs displayed high correlation with presupporting cells. In contrast, SLCs showed low transcriptional similarities to fetal Leydig cells, despite their known steroidogenic potential. Given that SLCs were more abundant in early rabbit ovaries than in testis, we performed this analysis in female and male datasets independently (Supp Fig. 7a-b). Again, SLCs displayed a high transcriptional correlation with presupporting cells, similar to what has been described in the mouse^39^, although the level of correlation was less prominent in males. In this dataset, SLCs also displayed low correlation with Leydig cells, indicating a distinct transcriptional identity (Supp Fig. 7a). We next explored whether the female enrichment of SLCs during early gonadogenesis corresponded with the activation of a sex-specific genetic program. For this purpose, we calculated the number of differentially expressed genes (DEG) between female and male SLCs and compared it with cell proportions across developmental stages. A progressive increase in upregulated genes was observed throughout the female stages peaking at 19 *dpc*. Importantly, this expression dynamic was concomitant with the increased proportion of female/male SLCs, suggesting a potential association between these processes (Fig 4b). The GO analysis on female SLCs DEGs at 19 *dpc* revealed the presence of terms related to transport of molecules, metabolic processes and post-translational modifications (Supp Fig. 8), suggesting a higher metabolic activity in female SLCs.

**Figure 7.**
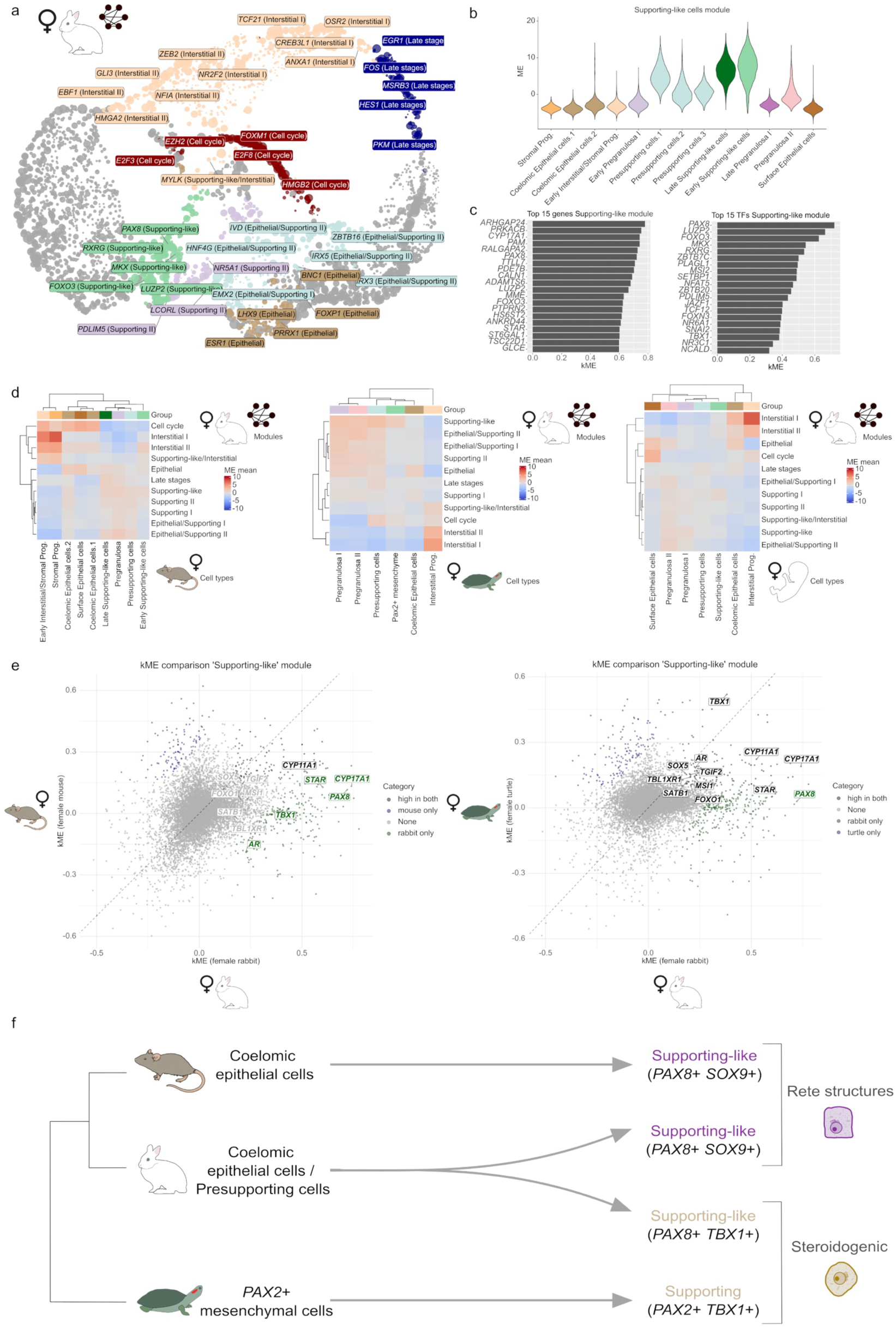
Activity of rabbit SLC network across species. a) UMAP projection of the female rabbit gene network computed with hdWGCNA. Genes are grouped based on their WGCNA module. The names for some top TFs per module are highlighted. b) Violin plot showing the Module Eigengene (ME) score per cell type of a module highly scored in female rabbit SLCs. This implies that the genes belonging to the module are predominantly expressed in those cells. c) Barplot showing the kMEs for the top 15 genes and the top 15 transcription factors of the Supporting-like module. The higher the kME the more the expression of a gene correlates with the general expression pattern of a module. d) Heatmap showing the ME scores of the rabbit WGCNA modules in mouse cell types (left panel), turtle cell types (middle panel) and human cell types (right panel). e) Scatterplot displaying the kMEs of the genes belonging to the rabbit SLC module in female mouse vs female rabbit (left panel) and female turtle vs female rabbit (right panel). Outliers were coloured using an IQR-based threshold. Highlighted are the androgen-synthesis genes, *PAX8* and transcription factors highly scored in rabbit and turtle but lowly scored in mouse and human. Green marks highlighted genes that are highly scored in the rabbit, black marks those that are highly scored in both species and grey marks those that do not surpass the threshold. f) Schematic representation of the origin of cell types and their similarities across species.

Given the transcriptional similarities between SLCs and the supporting lineage, and the dimorphic nature of the latter, we investigated whether such sex-specific patterns are also observed in SLCs. We trained a LASSO model to determine a score based on the transcriptional profile of pregranulosa I, pregranulosa II and Sertoli cells^39^. Then we used the models to compute these scores for different rabbit cell types including the early and late SLCs of the female and male gonads. As an internal control, pregranulosa II cells showed a high pregranulosa II score and a low Sertoli score (Fig 4c), and the opposite was observed for the Sertoli cells (Fig 4d). Similar to what was described in the mouse, female late SLCs exhibited higher pregranulosa II score than Sertoli score (Fig 4c) and the opposite was observed in the male counterpart (Fig 4d). Although more similarities were observed for male late SLCs with Sertoli than with pregranulosa cells, the similarity scores were generally low. This may be explained by the fact that our atlas contains two additional late stages for ovaries than for testes (stages 24 and 28). Thus, male late SLCs may represent a less differentiated state than female late SLCs. Intriguingly, the comparison of the pregranulosa I and pregranulosa II scores for the female late SLCs revealed a higher pregranulosa II-like signature (Supp Fig. 9a). Otherwise, no remarkable tendencies were observed for female or male early SLCs, consistent with their undifferentiated state. Next, we checked whether the male SLCs display a more similar transcriptional profile to that of the Sertoli cells than to the Fetal Leydig cell transcriptome, as Person correlation analyses previously suggested (Supp Fig. 7a). We trained another LASSO model to learn the molecular signature of Fetal Leydig cells and then compared the Sertoli score and the Leydig score for the male late SLCs. This analysis revealed that the male SLCs had a higher Sertoli than Leydig score, although the scores were again low (Fig 4d). When applied to female SLCs, these cells exhibited higher pregranulosa II than Leydig score (Fig 4c), indicating more similarity with the supporting than with the interstitial lineage.

We further explored the transcriptional similarities between rabbit SLCs and the supporting lineage, by examining differentially expressed genes at early and late developmental points. The number of DEGs between the female late SLCs and the two late supporting populations, pregranulosa II and late pregranulosa I, were substantially different. While 681 genes were upregulated in SLCs compared with pregranulosa II cells, up to 1734 genes were upregulated in SLCs in the comparison against late pregranulosa I cells (Supp Fig. 10a). These results, together with the results obtained with the LASSO models, suggest that female SLCs are more closely related to the pregranulosa II subpopulation. We subsequently performed a GO analysis on female SLCs upregulated genes and observed an enrichment for terms including “positive regulation of hormone metabolic process” or “C21-steroid hormone metabolic process” (Supp Fig. 11). Notably, we observed terms associated with “mesenchymal to epithelial transition involved in metanephros morphogenesis”, supporting transcriptional similarities with the mesonephros, as also described in the mouse^39^. In contrast, the equivalent analyses for male late SLCs and Sertoli cells revealed an enrichment for SLC-upregulated genes in GO terms like “metanephros morphogenesis”. Yet, the AUCell score of GO terms related to androgen synthesis, such as “regulation of steroid metabolic process” showed less specificity for the supporting-like lineage than in the female (Supp Fig. 11), consistent with the progressive shutdown of the steroidogenic pathway in male SLCs. Overall, our results demonstrate that, despite activating a sex hormone program, SLCs are more similar to the supporting than to the steroidogenic lineage. Notably, SLCs acquire a sexually dimorphic identity during sex determination that also extends to their steroidogenic activity.

### Trajectory analyses reveal the functional specialization of two distinct SLC subpopulations

To explore the developmental dynamics of rabbit SLCs, we compared them to their homologous counterparts in the mouse gonad, by using publicly available scRNA-seq datasets of analogous timepoints^39^. As shown above, cell-type proportion analysis revealed a much higher abundance of SLCs in rabbit gonads than in mice, pointing to species-specific differences in population dynamics (Fig 2a-d). We assessed the relationships of SLCs with other gonadal clusters by reconstructing a connectivity graph of the mouse and rabbit somatic gonads using PAGA^63^. Besides displaying a high degree of connectivity with late SLCs, mouse early SLCs of both sexes were predominantly connected with coelomic epithelial cells than with other cell types (Fig 5a-b, Supp Fig. 12), as previously described^39^. In contrast, the PAGA reconstruction revealed the existence of two distinct early SLC subpopulations in female rabbit gonads. The early SLC subcluster I (cluster 3) was strongly connected to presupporting cells (Fig 5c-d), displaying the high expression levels of the androgen synthesis genes and low levels of *WNT6* (Supp Fig. 13a). The early SLC subcluster II (cluster 7) was mostly connected to the coelomic epithelial cells (Fig 5c and e), showing less expression of androgen synthesis genes and almost no detectable expression of *WNT6* (Supp Fig. 13a). While these 2 subclusters were also present in the male rabbit gonad, the connection with the presupporting lineage was less evident (Supp Fig. 14). Of note, we also observed a degree of connectivity between early SLCs and presupporting cells in male mouse gonads, but not female, as previously described^39^ (Supp Fig. 12b-d). Overall, these results suggest that rabbit SLCs may be ontogenetically related to the coelomic epithelium, as also described in the mouse. In addition, a potential relationship with the presupporting lineage appears to exist specifically in the rabbit ovary. Yet, the nature of such a relationship is difficult to ascertain, as these two cell types coexist at all analyzed stages.

To further explore gene expression dynamics along the two potential differentiation trajectories of female rabbit SLCs, we selected clusters of cells strongly connected between them and to each subcluster. We subsequently computed the diffusion maps of those subsets, obtaining differentiating trajectories going from the coelomic epithelial or presupporting cells to the SLCs subclusters (Fig 6a, Supp Fig. 15a). Then, we employed Slingshot^64^ to compute a pseudotime value for each cell along the two different lineages and performed dynamic time warping to compare gene expression dynamics. Independently of the cellular origin, we observe that one of the terminal states upregulated and retained expression of the steroidogenic genes, whereas the other state did not (Fig 6b, Supp Fig. 15a-c). Indeed, cells from the non-steroidogenic population expressed *SOX9* and *KRT8* (Fig 6b, Supp Fig. 15c-d), all known markers of the rete ovarii^39^, suggesting a differentiation towards such lineage. Similar differentiation dynamics, although to a lesser extent, were observed in male rabbits (Supp Fig. 16a-f), but with lower expression of androgen-synthesis genes at later stages. More specifically, we observe the differentiation into two lineages, one of them retaining expression of steroid-synthesis genes and the other expressing Sertoli-like markers such as *AARD*, *SOX9*, *DMRT1* and higher expression of *KRT8*, suggesting that they might differentiate as components of the rete testis (Supp Fig. 16c and f). When performing equivalent analyses on female mouse data, we observed less clear terminal states for the SLCs than in the female rabbit ovary, with few informative genes showing different expression patterns. *PAX8* and *SOX9* were slightly more highly expressed in one of the terminal states, suggesting that those cells might commit towards rete ovarii formation, but no clear expression pattern was observed in the other potential terminal state. More importantly, and as expected, no remarkable expression pattern for the steroidogenic genes along the differentiation of SLCs was observed (Supp Fig. 17a-d).

Further 3D gonadal reconstructions and immunohistochemistry analyses revealed that in XY rabbit prenatal gonads, PAX8+ cells localized at SOX9-regions at the anterior pole, corresponding to the developing rete testis, whereas SOX9 expression was restricted to Sertoli cells within the seminiferous cords (Supp Fig. 18a–b). After birth, at 25 days *post-partum* (*dpp*), PAX8+ cells were detected in the testis medulla and contributed to the rete testis and Sertoli valve, where they co-expressed SOX9 (Supp Fig. 18c). Co-localization of PAX8+ and SOX9+ cells could also be observed in the prenatal XX gonad near the border with the mesonephros (Fig 6c and Supp Fig. 18d). However, these double-positive cells do not colocalize with the expression of the steroidogenic marker *CYP17A1*, which was instead restricted to a PAX8+ SOX9-population located in the medulla. Collectively, our findings suggest that rabbit SLCs have a dual potential, giving rise not only to rete precursors, as also described for mouse and human^39,40^, but also to a distinct steroid-producing population.

### Steroidogenic SLCs are transcriptionally similar to non-mammalian PAX+ mesenchymal cells

It was recently shown that steroid-producing cells derived from the supporting lineage are present during early gonadogenesis in non-mammalian vertebrates^41,42^. To explore whether rabbit SLCs may be ontologically related to these steroidogenic supporting cells, we reconstructed and compared coexpression gene modules using hdWGCNA. For this purpose, we made use of a publicly available scRNA-seq dataset from turtle gonads^41^. We also included data from mammalian species like mouse and human in our comparison. Humans are particularly interesting, as it was reported that early gonads from this species can also synthesize estrogens during prenatal gonadal development^65,66^. After processing all datasets with the same workflows, we used hdWGCNA^67^ to infer modules of coexpressed genes. To capture sex-specific transcriptional networks, the analysis was performed separately for female and male samples, where SLCs retain or lack steroidogenic activity, respectively. Coexpression modules were subsequently annotated based on the cell type in which they displayed the highest activity. For the female rabbit dataset, we obtained 11 specific modules (Fig 7a), one of which showed high activity in SLCs and its precursor population, the presupporting cells (Fig 7b). We subsequently explored the genes displaying the highest kME, which measures how tightly a gene follows the overall expression pattern of the module. Among those, we found steroid-related genes such as *CYP17A1* or *STAR* (Fig 7c). The top 15 transcription factors (TFs) associated with this module included *PAX8*, but also other factors involved in lineage differentiation like *RXRG, MKX*, *PLAGL1* or *TBX1* (Fig 7c). As observed in differentiation trajectory analyses (Fig 6a-b), *TBX1* was highly expressed in the SLC lineage that retains the expression of the androgen synthesis genes. Interestingly, *TBX1* was also expressed in the turtle supporting lineage, while not being expressed in the SLCs of mice or lowly expressed in that of humans (Supp Fig. 19). To explore the cross-species activity of rabbit gene modules, we projected them onto the datasets of the other species and calculated Module Eigengene (ME) values, a measure that reflects the cells in which the genes of each module are predominantly expressed. Modules associated with interstitial, Sertoli or fetal Leydig cells display a high degree of activity in the corresponding cell types of other species (Fig 7d and Supp Fig. 20a-c). In females, the rabbit SLC module displays activity in the SLCs of other mammals, but also in pregranulosa cells, confirming the transcriptional similarities observed previously (Fig 4a and c). Remarkably, the ME scores for the rabbit SLC module indicate activity in with the supporting lineage of the turtle, despite the large evolutionary distance separating both species (Fig 7d). Importantly, the turtle supporting lineage was recently shown to express *TBX1* and to be steroidogenically active^41^. Notably, the rabbit SLC module was the only one that also showed activity in the turtle *Pax2+* mesenchymal population, the precursors of steroidogenic supporting cells in this species, which express *Pax2*, a functionally redundant paralog of *PAX8*.

To explore the conservation of expression for the individual genes of the rabbit SLCs module, we compared their predicted kMEs across species (Fig 7e). These analyses confirmed the high kME values of the steroid-synthesis genes mainly in the rabbit and in the turtle, while remaining low in the mouse. Among the genes that showed high kME values across all the species in the SLCs module we found *AMHR2* and *NR5A1* (Supp Fig. 21a-c). Interestingly, besides the steroid-synthesis genes, the transcription factor *TBX1* exhibited high kME values in both rabbit and turtle, confirming that the projection of the module was done primarily to the steroidogenic cells of the turtle. For the male rabbit dataset we obtained 10 specific modules, two of which were specific to SLCs (Supp Fig. 22a-b). Further inspection of the SLC module with the highest ME score in the SLCs revealed the presence of *PAX8* and *RXRG*, but no gene related to steroid synthesis was detected (Supp Fig. 22d-e). The projection of the SLCs module with the highest activity to the other species revealed a similar pattern to that of the female, with its activity being generalized in the supporting lineage (Supp Fig. 20). Interestingly, the projected module also exhibited medium ME score in the fetal Leydig cells of both human and mouse, predominantly in the mouse (Supp Fig. 20a-c).

To confirm the potential homology between the turtle *PAX2*+ cells and the rabbit *PAX8*+ cells, we ran hdWGCNA in the turtle data and projected the modules into the rabbit (Supp Fig. 23). We obtained a module highly specific of the *PAX2+* mesenchymal cells (Supp Fig. 23d). The projection of this turtle *PAX2*+ mesenchymal module showed more activity in the rabbit SLCs (Supp Fig. 23c-e), suggesting the equivalence between both cell types. Among the genes with a high kME predominantly in the turtle we found *PAX2*, and in the rabbit *PAX8*, as expected, whereas *TBX1* was highly scored in both (Supp Fig. 23b). To further validate the similarities between rabbit SLCs and turtle *PAX2*+ mesenchymal cells as putative homologous cell types, we performed an unsupervised MetaNeighbor analysis^68^. As expected, cells from the same organism tended to yield higher AUROC scores, reflecting species-specific signatures. However, rabbit pre-supporting and SLCs achieved AUROCs above 0.5 compared with turtle presupporting and *PAX2⁺* mesenchymal cells (Supp Fig. 24a-d). An AUROC value of 0.5 indicates similarities at the level of random expectations, therefore values higher than 0.5 indicate a degree of transcriptional similarity between cell types. Altogether, our results suggest an evolutionary relationship between the SLCs of the rabbit and the steroidogenic supporting lineage of the turtle, both deriving from PAX+ precursors and expressing lineage-specific transcription factors, such as *TBX1* (Fig 7f).

## DISCUSSION

Studies on sexual development have predominantly focused on characterizing a limited number of gonadal cell lineages. These mostly include the germinal lineage, which undergoes gametogenesis; the supporting lineage, which initiates sex determination and the interstitial lineage, which provides steroidogenic activity. While this framework has been very influential, recent single-cell studies suggest that it may oversimplify the complexity and plasticity of gonadal cell fates^39,42,41,40^. Here we demonstrate that an alternative lineage, the SLCs, can perform steroidogenic functions that are analogous to that of interstitially derived cells. SLCs were recently identified as an independent lineage through single-cell analyses of mouse gonads, emerging as early as E10.5 and following a distinct developmental trajectory to other cell types^39^. Their elusive nature and late discovery can be partly attributed to their low abundance in gonads from mice, or from other species like humans. Functionally, SLCs have been shown to have a structural function, contributing to the rete ovarii and rete testis, which are essential for gamete transport^39,69,70^. In the rabbit, we show that SLCs not only perform this structural function, but also support early steroid production, a key process for ovarian development and folliculogenesis^27^. In contrast to other well-studied mammalian models, SLCs constitute a considerable percentage of the rabbit early gonad, particularly in ovaries. It is worth noting that the steroidogenic program is also initiated in male rabbit SLCs but progressively substituted by the activity of Leydig cells. This may suggest the existence of a regulatory crosstalk between these two cell types, as the expression of steroidogenic genes declines in SLCs while increasing in Leydig cells. This decline in steroid-related gene expression occurs in the presence of DHH, a potent morphogen that promotes the differentiation of interstitial progenitors into steroidogenic theca or fetal Leydig cells^7^, suggesting that the analogous activity in SLCs involves alternative pathways to Hedgehog signaling. Taken together, our findings challenge the long-standing paradigm that gonadal steroidogenesis requires the action of interstitially derived cells.

While the steroidogenic potential of mammalian interstitial lineages is undisputed, comparative studies in non-mammalian vertebrates further indicate the existence of alternative cellular sources for sex hormones. In the turtle *Trachemys scripta*, a species where steroids play a critical role in sex determination, the interstitial lineage does not exhibit steroidogenic activity, as denoted by the absence of Leydig cells in early gonads. Instead, this pathway is active in the supporting lineage and its derivatives, thus highlighting their intrinsic steroidogenic potential^41^. Similarly, the supporting cells of the chicken gonad give rise to steroidogenic cells during early gonadogenesis^42^. These non-mammalian examples provide an important anchor point for inferring the evolutionary origin of the steroidogenic activity of mammalian SLCs. Despite the vast evolutionary distances, our comparative analyses reveal common transcriptional signatures between the steroidogenic supporting cells of turtles and rabbit SLCs. Both lineages express key specification markers like *PAX2 or PAX8,* which are paralogous genes with redundant function, and *TBX1*. In addition, these two lineages occupy similar anatomical positions at the border between gonad and mesonephros, suggesting that they may represent homologous cell types. Collectively, these findings delineate two evolutionary scenarios that are non-mutually exclusive. One possibility is that the gonads of the amniotes common ancestor relied on supporting-derived steroidogenic cells as a primary source of steroids, with a subsequent specialization of fetal interstitial steroidogenic cells in mammals. Alternatively, supporting lineages may retain a latent steroidogenic potential that could be transiently activated at specific developmental windows or in certain taxa. In fact, in some species like the mouse, fetal Sertoli cells can also express *HSD17B3*^49^, which encodes an enzyme that contributes to the conversion of androstenedione to testosterone. In the mammalian postnatal ovary, granulosa cells are also known to facilitate the conversion of the androgens produced by theca cells into estrogens^71,72^. Further evidence supporting this latent potential comes from perturbation experiments in mice and chicken showing that the disruption of factors like WT1 or TOX3 can steer the supporting lineage to ectopically activate the full steroidogenic program^73,74^. Moreover, in *FOXL2* knockout XX goats, which display female-to-male sex reversal, the steroidogenic *CYP11A1* and *HSD17B3* genes are upregulated prior to the complete differentiation of Leydig cells^75^. Altogether, these observations highlight the inherent plasticity of the gonadal lineages, despite the overall conservation of functions across vertebrate species.

The coordinated activation of steroidogenic genes observed in both SLCs and Leydig cells suggests a shared gene regulatory program across these lineages. This is supported by the expression of *NR5A1*, a known activator of steroidogenesis, in both cell types. Yet, we also observe evidence of cell-type specific differences. For example, female rabbit SLCs can convert pregnenolone to progesterone in the absence of *HSD3B* expression. In contrast, this gene is prominently active in the steroidogenic program of male Leydig cells. Our screening for genes that may compensate for the absence of *HSD3B* expression in SLCs pinpointed *PGER5* as a potential candidate, due to its temporal overlap with the steroidogenic window. *PGER5* belongs to the *AKR1C* gene family^76^, which has been shown to be capable of performing the 3β-hydroxysteroid dehydrogenase function^62^. This is consistent with the low levels of 3β-hydroxysteroid dehydrogenase activity previously reported in the medullary region of fetal rabbit ovaries, the anatomical region where SLC are localized^77^. Although an additional isomerase activity is also required for the full conversion of pregnenolone to progesterone, our steroid profiling demonstrates that this step effectively occurs. Yet, if *PGER5* or other genes that may have not been annotated in the rabbit genome can perform such isomerase activity remains to be tested. Besides the potential compensatory effect of *PGER5*, we observe that Progesterone is detected at low levels and estrogens could not be detected in early rabbit ovaries, due to known technical limitations of LC-MS. Nevertheless, estrogens have been confirmed to be present in the prenatal rabbit ovary through alternative methods and their functional relevance demonstrated *in vivo*^28,30,31,27,29^. These studies, together with our results, may point to low endogenous estrogen levels that are likely exerting a local effect in the gonad, rather than broad systemic effects at the embryonic level. In fact, the concept of localized hormonal action was exemplified by the pioneering work of Alfred Jost, which demonstrated that the stabilization of the Wolffian ducts occurs through local androgen signaling in the embryonic rabbit testis^78,79^. In any case, our results again suggest the existence of distinct regulatory mechanisms underlying the activation of steroidogenesis in SLCs and interstitial cells.

The capacity of supporting-related lineages to assume steroidogenic roles also has important implications for our understanding of gonadal development, suggesting a dynamic allocation of functional roles during sex differentiation. Further evidence for this flexibility comes from studies in species with natural variations in steroidogenesis, like the iberian mole *Talpa occidentalis*. In this species, female individuals develop ovotestes that produce high levels of androgens that cause masculinization and increased muscular development^80,81^. In female ovotestes, the steroidogenic activity is observed in interstitial-like cells that do not express supporting cell markers such as FOXL2, but that are distinct from the lineage that differentiates into theca cell^82^. Moreover, single-cell atlases of human fetal ovaries, where steroidogenesis has also been reported, indicate that this activity is more diffuse across cell types than in mouse or rabbit gonads. Furthermore, SLC-like populations do exist but represent a minoritarian lineage in the human gonad. These observations further support the notion that the steroidogenic function of the developing gonad is not rigidly restricted to a single lineage, but can emerge from multiple cellular lineages, depending on developmental or evolutionary contexts.

Collectively, the findings described here redefine a classical paradigm by demonstrating that steroidogenic function in early mammalian gonads can emerge outside the canonical interstitial lineage. This inherent flexibility is also relevant for the study of Differences of Sex Development (DSDs), many of which cannot be explained solely by mutations in canonical sex-determining genes. It is conceivable that some unresolved cases may be explained by regulatory mutations that ectopically activate the steroidogenic function in cellular lineages that have a latent potential. In a broader context, our results highlight that vertebrate species may achieve similar reproductive outcomes through flexible cellular strategies rather than a strict allocation of functions.

## MATERIALS AND METHODS

### Gonad collections and dissociation

Pregnant New Zealand White rabbits at the stages 13-21 *dpc* were obtained from Charles River (France branch). For the latest developmental stages, sex was initially determined based on anatomical differences, particularly the presence of prominent vasculature in testes and its absence in ovaries. In all stages, sex was confirmed by PCR amplification of the Sry locus. We pooled 3-4 pairs of gonads per sex and stage. Dissected gonads were dissociated into single-cells by incubating them with 0.05% Trypsin-EDTA (Gibco cat#25300-054) in addition to 1/10 portion of 5% BSA (Sigma-Aldrich cat#A7030-50G) /DPBS for 7-10 minutes at 37°C. Dissociation was facilitated by pipetting gently every 2-3 minutes. Residual connective tissue, which appeared as whitish clumps at the end of the dissociation process, was discarded. Trypsin activity was halted by adding 2 portions of 5% BSA/DPBS. Cell suspension was passed through a 40 µm FlowMi cell strainer (Fisher Scientific cat#15342931) to remove cell clumps and debris. Filtrated cells were collected by centrifugation at 250-300g for 5 minutes at 4°C. The cell pellet was resuspended in DPBS for counting and viability check using 50% v/v trypan blue staining (Sigma Aldrich cat#T8154-20ML) under a light microscope. Only samples with ≥ 80% were used for downstream single-cell library preparation. Finally, cells were fixed with 98% methanol added drop by drop and stored in −80°C freezer.

For 24 and 28 *dpc*, fetal ovaries were obtained from New Zealand rabbits bred at the SAJ rabbit facility (Jouy-en-Josas, France). The two gonads from the same fetus were pooled per stage in 500 µl of Trypsin-EDTA 0.05% at 37°C for 30 min. Dissociation was facilitated by gently pipetting every 5 minutes. Trypsin activity was stopped by adding SVF for a final 2% concentration. The cell suspension was passed through a 70 µm filter, then a 40 µm FlowMi Cell Strainer (Sigma-Aldrich cat #BAH136800040) to remove cell clumps and debris. The filtrated cells were collected by centrifugation at 300g for 10 minutes. The cell pellet was resuspended in 500 µL (24 *dpc*) or 1 mL (28 *dpc*) of DMEM/F12-SVF 2%. To determine whether the obtained cells were suitable for downstream analysis (cell viability > 80%), cell viability was evaluated using trypan blue staining with a haemocytometer r (TC20, Bio-Rad cat#1450102). The cell concentration was then adjusted to around 1000 cells/μl before loading onto the single-cell chip.

### 10x Genomics library preparation and sequencing

For the data from gonads at the stages 13-21 *dpc,* fixed cells were equilibrated to 4°C on ice for at least 5 minutes. Subsequently, the cells were pelleted at 1,000g for 5 minutes at 4℃. The cells were rehydrated and gently washed with rehydration buffer (0.04% BSA, 1mM DTT (Life Technologies cat#707265ML), 0.4U/µl RNaseIn (Life Technologies cat#AM2682), 0.2U/µl SuperaseIn (Life Technologies cat#AM2696), in DPBS), using wide-bore tips (Corning Axygen Cat#T-205-WB-C-R-S). Cell number and quality after rehydration were assessed using 50% v/v trypan blue staining. Cell suspensions were adjusted to a final concentration of 700-1,200 cells/µl. Single-cell libraries were generated using the Chromium Next GEM Single Cell 3ʹ Kit v3.1 (10X Genomics), according to manufacturer’s instructions, targeting 10,000 cells per reaction. Two libraries were prepared for each developmental stage and sex, except for female gonads at stage 15, for which only one library was prepared. Libraries were sequenced to a minimum depth of 200 million reads per library on the Illumina Novaseq 6000 or DNBSEQ-G400 systems using the paired-end sequencing format: Read 1, 28 cycles; I7 index, 10 cycles; I5 index, 10 cycles; Read 2, 90 cycles.

For the data from ovaries at 24 and 28 *dpc*, Gel-Bead in Emulsions (GEMs) were generated using a Chromium 10X Single Cell System (10X Genomics) and cDNA libraries were prepared using the Chromium Next GEM Single Cell 5’ Reagent v2 (10X Genomics). Two libraries were prepared for each developmental stage. The I2BC High-throughput Sequencing Platform (https://www.i2bc.paris-saclay.fr/sequencing/ng-sequencing/; Université Paris-Saclay, Gif-sur-Yvette, France) provided the facilities and expertise for library preparation and sequencing (Paired-end 28-64 bp, Illumina NextSeq500).

### Alignment of reads, preprocessing and integration

The sequenced scRNA-seq libraries were processed with 10X Genomics Cell Ranger v7.0.2 using our optimized *Oryctolagus cuniculus* reference genome and with the default parameters. The genome annotation was improved by correcting and extending the three prime ends of the genes. The filtered count matrix of each library returned by Cell Ranger was loaded into Scanpy^83^, and the pipeline with the recommended standard practices was followed. As a first filter, cells expressing fewer than 200 genes and genes expressed in less than 3 cells were excluded from the analysis. We also removed cells that show more than 5 median absolute deviations (MADs) from the median of the total counts or from the median of the number of detected genes. Cells with over 10% mitochondrial reads (5% for the mouse and 20% for the human and turtle datasets) were also removed. Doublets were detected individually for each library using Scrublet^84^ with a value of 0.15 (0.1 for the human data) for the parameter expected_doublet_rate to detect the expected 5% of doublets per library. Cells with a doublet score equal or higher to 0.3 were later removed from the analysis. To detect droplets containing damaged cells, we used DropletQC^85^ and clusters of cells with a high nuclear fraction score and low UMI counts were later excluded. After the normalization and log-transformation of the raw counts with the default Scanpy settings, we computed the top 5000 highly variable genes on a library basis, and the data was scaled to a maximum value of 10. The cell cycle scores were calculated using the score_genes_cell_cycle of Scanpy based on the list of S genes and G2M genes provided by Seurat^86^. Then, using scVI^87^, we corrected for batch effects using as covariate keys the S and G2M cell cycle scores. The correction for batch effects was done for the full dataset and for the somatic compartment on a sex basis. Using the integrated latent representation, we computed the neighbour graph, the Leiden clustering and the UMAP visualization with Scanpy functions. Clusters were annotated based on well-known mouse markers using the 1.4 Leiden resolution. To achieve a higher resolution in the annotation, the clustering and annotation was then refined for the clusters identified as part of the somatic gonad for the female and male separately. Differential abundance analysis was conducted in the full dataset using milo^88^, available in the pertpy^89^ package, following the recommendations of the vignette. Correlations between transcriptomes were calculated using Pearson on the mean expression of genes by cell type. The hierarchical clustering used for plotting the matrix of correlations was then obtained with the clustermap function of the seaborn package.

### Mouse, human and turtle scRNA-seq datasets

Unless specified another way, mouse, human and turtle datasets^39,40,41^ were processed as previously described for the rabbit dataset. The annotation of cells was based on the original annotations provided by the authors, being adapted to match the clustering and standardized to the annotation nomenclature used for the rabbit.

### Protein sequence similarity

The protein sequence alignment between rabbit PGER5 and human AKR1C3 proteins was performed using the R package msa^90^ (Multiple Sequence Alignment) with the ClustalOmega algorithm.

### Lineage reconstruction and dynamic-time-wrapping

The lineage reconstruction of the somatic gonadal clusters was conducted using the Partition-based graph abstraction (PAGA^63^) with the default parameters based on the neighbour graph (n_neighbors = 15) computed from the scVI components. An additional subclustering was performed for some coelomic epithelium and SLCs clusters to obtain cleaner connections between finer clusters. PAGA estimates connectivity between clusters and provides a graph-like map that preserves the global topology of the data. The embedding for the somatic gonadal cells was then initialized using PAGA-initialization by specifying the parameter init_pos as “paga” in the Scanpy umap function. Clusters most strongly linked to early supporting-like cells were chosen for diffusion-map analysis, ensuring coverage from early to late developmental stages. Diffusion maps were computed from a k-nearest-neighbors graph built with Scanpy’s neighbors function (n_neighbors = 10, n_pcs = 15) in the scVI latent space. Using the resulting diffusion components, we inferred differentiation trajectories with the slingshot^64^ package and assigned to each cell a pseudotime value. In the getLineages function, start.clus was set to the clusters corresponding to the earliest developmental points and end.clus to the clusters corresponding to the latest stages. We analysed gene-expression dynamics along the inferred trajectories using the dtwclust^91^ package to group genes with similar expression trends over pseudotime. Cells were binned into 20 pseudotime intervals, and for each gene the mean expression within each interval was computed. We then performed temporal-series clustering with tsclust (type = “partitional”, k = 5).

### Differential expression analysis and GO term enrichment analysis

The differential expressed genes were computed using the differential_expression function of scvi_tools, comparing no more than two cell types at a time. The markers were then filtered to only keep those with a false discovery rate (FDR) lower than 0.1 and a logFC greater than 0. The GO enrichment analysis was then computed using the function compareCluster from the clusterProfiler^92^ package with the *mus musculus* database as Orgdb, the fun parameter set to enrichGO and Biological Processes as GO category. Similar GO terms were collapsed using the GOSemSim^93^ package with an h of 0.5 in the cutree function. Sets of genes of interest were scored using AUCell^94^.

### Lasso classifiers

To assess the molecular signature of the different cell types, we randomly selected 600 cells per annotated cell type and we trained a LASSO classifier using the LassoCV function from the scikit-learn^95^ module. Through L1 regularization, the LASSO framework identifies a subset of genes that define the transcriptional profile of the cells of interest. Several cross-validations folds were tried to ensure the correct training of the model and the max_iter parameter was set to 10,000.

### Cross-species analysis

For the cross-species comparisons the gene identifiers of the different species were converted to human SYMBOL using the list of orthologues provided by the NCBI. Only genes with 1 to 1 mapping were conserved. For the hdWGCNA^67^ analysis, the AnnData objects processed with Scanpy and encompassing only somatic gonad cell types were loaded into the Seurat workspace by using the SeuratDisk package. These Seurat objects were then filtered to only keep the subset of common orthologue genes, and they were prepared for the hdWGCNA analysis following the steps of the tutorial. In the MetacellByGroups function, metacells were created based on their sample of origin and the cell type annotation, and the parameter k and max_shared were set to 25 and 10, respectively. The soft powers for the ConstructNetwork function were manually selected based on the visual inspection of the plots and the minModuleSize was set to 20. The module names were renamed based on the cell types in which they exhibit the highest activity. The networks were generated on a sex and species basis. Once generated, the female and male rabbit networks were projected into the other species by using the rabbit dataset as reference and the other species dataset as a query in the ProjectModules function. The projection of the turtle modules to the rabbit dataset was done using the same approach, but no splitting by sex was conducted. For visualization in scatter plots, the differences between the kMEs of the two species were computed and standardized using Z-scores. Then, outliers were detected using an IQR-based threshold.

To explore cell type similarities across different species we ran the unsupervised version of Metaneighbor^96^ on the orthologous gene set on a sex basis. Before running it, the top 10.000 variable features of each species were selected and the intersection was computed. Due to the high number of samples and genes, the fast version of MetaNeighborUS was used.

### Steroidomics

Multi-targeted extended steroid profiling of male and female rabbit fetal gonads was performed following a previously published protocol^59^ applied to 173 targeted metabolites (Supp. Table 1). Within the steroid profile, nine compounds could be accurately quantified with a one-point internal calibration workflow: cortisol (F), cortisone (E), 11-deoxycortisol (S), pregnenolone (P5), progesterone (P4), 17α-hydroxyprogesterone (17α-OHP4), androstenedione (A4), testosterone (T), and 5α-dihydrotestosterone (DHT)^59^. Twenty milligrams of fetal gonad tissue were homogenized in acetonitrile: methanol (9:1), using Beadbug-prefilled tubes and a Precellys instrument (Bertin Technologies). Extracts were filtered through HLB Prime cartridges (Waters Corp.), evaporated to dryness and reconstituted in methanol: water (1:1) before liquid chromatography – tandem mass spectrometry (LC-MS/MS) analysis. LC separation of steroids was performed with a biphenyl stationary phase (Restek Raptor Inert Biphenyl, 2.1 x 100 mm, 1.8 µm), and water and methanol with 0.01% formic acid were used as mobile phases. MS/MS data were acquired in Multiple Reaction Monitoring (MRM) with positive / negative polarity switching on a Triple Quadrupole instrument (Xevo TQ-XS, Waters Corp.). Peak areas and absolute concentrations were normalized by the exact mass of tissue. Both multivariate (Principal Component Analysis (PCA) and Orthogonal Partial Least Square – Discriminant Analysis (OPLS-DA)) and univariate analyses (two-tailed t-tests) were performed in SIMCA (Version 17.0.2, Sartorius AG) and Prism (Version 10.3.1, GraphPad), respectively.

### Tissue processing for whole-mount immunofluorescence and in-toto image acquisition of rabbit gonads

Whole-mount immunofluorescence followed by ethyl-cinnamate tissue clearing was performed as previously described for adult mouse ovaries^97^. Briefly, ovaries from 16, 24, and 28 *dpc* male and female rabbits were dissected in PBS (Day 0), fixed for 48-96 hours at room 4°C in 4% PFA/PBS, and gradually dehydrated in 100% methanol for storage at −20°C. When ready for analysis (Day 1), samples were progressively rehydrated to PBS 0.2% Triton X-100 (PTx.2) following an isopropanol gradient: 70% isopropanol / PBS; 50% isopropanol / PBS; 30% isopropanol / PBS; 100% PTx.2. Samples were then permeabilized overnight at 37°C in PTx.2 with 2.3% glycine. On day 2, samples were incubated in blocking solution (PTx.2 10% horse serum solution) for 6 h at 37°C. Tissues were then incubated in primary antibodies diluted in PTx.2 with 0.001% heparin (PTwH) with 10% horse serum for 72h at 37 °C. On day 5, samples were washed 3 times for 1 hour at RT in PTwH followed by incubation at 37°C for 48h with secondary antibodies diluted in PTwH with 10% horse serum. On day 7, after washing 3 times for 1 hour in PTwH, samples were progressively dehydrated into 100% isopropanol (30% isopropanol / PBS; 50% isopropanol / PBS; 70% isopropanol / PBS; 100% isopropanol) and left in 100% isopropanol overnight. On Day 8, samples were cleared in ethyl cinnamate (ECi). Samples were left overnight in ECi to allow for sufficient clearing prior to imaging. Antibodies used were: Rabbit anti-PAX8 (10226-1-AP, Proteintech); Rabbit anti-SOX9 (AB5535, Millipore-Sigma); Rabbit anti-CYP17A1 (14447-1-AP, Proteintech); Mouse anti-DMRT1 (sc-377167, Santa Cruz Biotechnologies); Goat anti-SOX9 (AF3076, R&D Systems); Rabbit ATTO-647 anti-mouse IgG (610-456-C46S, Rockland); Cy3 Donkey anti-goat (705-165-147, Jackson ImmunoResearch). Rabbit primary antibodies were conjugated using the Proteintech Flexable 2.0 AF647- and AF750-conjugation kits.

ECi-cleared ovaries were mounted in a 3D-printed coverslip holder (model available at https://www.mckeylab.com/3d-prints) as previously described for adult mouse ovaries^97^. Images were acquired using an Andor Dragonfly (Oxford Instruments, Abingdon, UK) with a Leica DMi8 microscope stand (Leica Microsystems, Wetzlar, Germany). Z-stacks were acquired with a 3µm interval using a 10x dry objective. Images were processed using Imaris software (Andor Technology, Belfast, UK) to generate figure displays and 3D segmentations (using the surfaces module). Processed images were assembled into figures using Adobe Photoshop.

### Immunohistochemistry analyses

Immediately after sampling, gonads were immersed in 4% paraformaldehyde (PFA) in phosphate-buffered saline (PBS). After 48 hours of fixation at 4°C, tissues were washed three times with PBS and stored at 4°C in 70% ethanol until paraffin inclusions. Adjacent sections of 5 µm thick were processed using a microtome (Leica RM2245) and organized on Superfrost Plus Slides (J18000AMNZ, Epredia). Before staining or experiments, sections were deparaffinized and rehydrated in successive baths of xylene and ethanol at room temperature.

Immunohistochemistry (IHC) was performed using the ABC amplification signal kit (PK-6100, Vector Laboratories) and DAB enzymatic reaction (SK-4100, Vector Laboratories). Briefly, the antigenic sites were unmasked with a citrate buffer (pH 6; H-3300, Vector Laboratories), and endogenous peroxidases were blocked with a 3% H2O2 solution (H1009, Sigma-Aldrich). Sections were then permeabilized with 1X PBS, 1% Bovine Serum Albumin (A7906, Sigma-Aldrich), and 0.2% Saponin (7395, Merck) and incubated overnight at 4°C with primary antibodies (rabbit anti-PAX8, 10226-1-AP, Proteintech; rabbit anti-SOX9, Francis Poulat; mouse anti-DMRT1, sc-377167, Santa Cruz; rabbit anti-FOXL2, homemade^98^). Following PBS washes, sections were incubated with biotinylated secondary antibodies (horse anti-rabbit, BA-1100, Vector Laboratories; anti-mouse, M.O.M kit BMK-2202, Vector Laboratories). After the ABC kit incubation and DAB revelation, hematoxylin staining was briefly performed to visualize the whole tissue. All stained sections were scanned using a 3DHISTECH panoramic scanner at the @Bridge platform (INRAE, Jouy-en-Josas, France).

## Supporting information

Barbera-Aura_et_al_Supplementary_material

## DATA AVAILABILITY

Single cell RNA-seq (scRNA-seq) data from rabbit gonads have been deposited in GEO repository (GSE331375). Equivalent single cell datasets from mouse, human and turtle are available at NCBI Gene Expression Omnibus GEO GSE184708, ArrayExpress with the accession number E-MTAB-10551 and GEO GSE271230, respectively.

## CODE AVAILABILITY

The custom R and python code to reproduce the analyses and the figures is available in GitHub (https://github.com/IvanBarbera00/scRNA_fetal_rabbit_gonad)

## AUTHOR CONTRIBUTIONS

I.B.A., W.Y.C., S.N., E.P., and D.G.L. conceived the study and designed experiments. W.Y.C., A.H.M., E.P., and E.D. collected and preprocessed rabbit gonads. W.Y.C., A.H.M. and Y.J. and A.H.M. performed single-cell RNA-seq experiments. J.M. and E.D. performed immunostainings. J.M. performed in toto gonad image acquisition and processing. M.Ga. performed and analyzed steroidomics experiments. I.B.A., W.Y.C., B.M.P. and E.D. analyzed data with inputs from R.D.A., C.M., M.Gu., S.N., E.P. and D.G.L. I.B.A. and D.G.L wrote the paper with input from all authors.

## ACKNOWLEDGEMENTS

We thank the animal facilities of the Max-Delbrück Center for Molecular Medicine and of the Universidad Pablo de Olavide for technical assistance. We thank S. Reissert-Oppermann, C. Westphal, J. López-Rios, J.M Delgado, A. Quetglas, J.M. Gonzalez and F.M. Real for their support with the animal work. We thank F.M. Real and members of the D.G.L. laboratory for their valuable input and comments on the paper. We thank P. Congar, G. Morin, and all the staff of the facility (SAJ, INRAE, Jouy-en-Josas, France) for the care of the rabbits; V. Gelin for monitoring rabbit pregnancies; J. Rivière and M. Vilotte (UMR GABI, INRAE, Jouy-en-Josas, France) for their assistance on the histological platform (@Bridge platform); M. Pannetier, M. André, A. Dewaele and E. Canon for their assistance on ovaries dissociation experiments; A. Frambourg and L. Jouneau for their assistance on bioinformatic analyses. We acknowledge the sequencing and bioinformatics expertise of the I2BC High-throughput sequencing facility, supported by France Génomique (funded by the French National Program “Investissement d’Avenir” ANR-10-INBS-09). Research in the Lupiañez lab was funded by the European Research Council (grant no. 101045439, 3D-REVOLUTION) and by the Spanish “Agencia Estatal de Investigación” (grant number PID2022-143253NB-I00/AEI/10.13039/501100011033/ FEDER, UE). Funded by the European Union. Views and opinions expressed are however those of the author(s) only and do not necessarily reflect those of the European Union or the European Research Council Executive Agency. Neither the European Union nor the granting authority can be held responsible for them. I.B. was supported by an FPU contract (Formación de Profesorado Universitario 2023 FPU23/00760) from the “Ministerio de Ciencia, Innovación y Universidades”. R.D.A. was supported by an EMBO Postdoctoral Fellowship (Grant no. EMBO ALTF 537-2020) and by the “Agencia Estatal de Investigación” (Ramón y Cajal RYC2023-045620-I). The SN lab was funded by the Swiss National Science Foundation, grants #31003A_173070, #310030_200316, #10.001.251. J.M. was supported by startup funds from the Department of Pediatrics at CU Anschutz and by the National Institutes of Health grant # R00HD103778 to JM. Research in the Pailhoux team was funded by the French ANR “Agence Nationale de la Recherche” grants (RNA-SEX: ANR-19-CE14-0012; ARDIGERM: ANR-20-CE14-0022) and the PHASE department of INRAE (API – CAROT-2019). E.D. was supported by the ANR RNA-SEX and the PHASE department of INRAE.

