## Supplementary material for "Supporting-like cells constitute an alternative steroidogenic lineage conserved in amniotes": Barbera-Aura_et_al_Supplementary_material

[1]*Centro Andaluz de Biología del Desarrollo (CABD), Consejo Superior de Investigaciones Científicas/Universidad Pablo de Olavide/Junta de Andalucía, Sevilla, Spain*
[2]*Max-Delbrück Center for Molecular Medicine in the Helmholtz Association (MDC), Berlin Institute for Medical Systems Biology (BIMSB), Epigenetics and Sex Development Group, Berlin, Germany*
[3]*Current address: Institute of Molecular and Clinical Ophthalmology, Basel, Switzerland*
[4]*Université Paris-Saclay, UVSQ, INRAE, BREED, 78350, Jouy-en-Josas, France*
[5]*École Nationale Vétérinaire d'Alfort, BREED, 94700, Maisons-Alfort, France*
[6]*Current address: Institute for Reproductive and Developmental Sciences, Department of Pathology and Laboratory Medicine, University of Kansas Medical Center, Kansas City, KS, USA*
[7]*School of Pharmaceutical Sciences, University of Geneva, CMU, Geneva, Switzerland*
[8]*Swiss Centre for Applied Human Toxicology (SCAHT), Basel, Switzerland*
[9]*Institute of Pharmaceutical Sciences of Western Switzerland, University of Geneva, CMU, Geneva, Switzerland*
[10]*Department of Genetic Medicine and Development, University of Geneva, Geneva, Switzerland*
[11]*Bioinformatics and Omics Data Science Platform, Max Delbruck Center for Molecular Medicine, The Berlin Institute for Molecular Systems Biology, Hannoversche Str. 28, 10115, Berlin, Germany*
[12]*Université Paris-Saclay, CEA, CNRS, Institute for Integrative Biology of the Cell (I2BC), Gif-sur-Yvette 91198, France.*
[13]*Section of Developmental Biology, Department of Pediatrics, University of Colorado Anschutz, Aurora, CO, USA*
[*]*These authors contributed equally: Iván Barberá-Aura, Wai-Yee Chung*

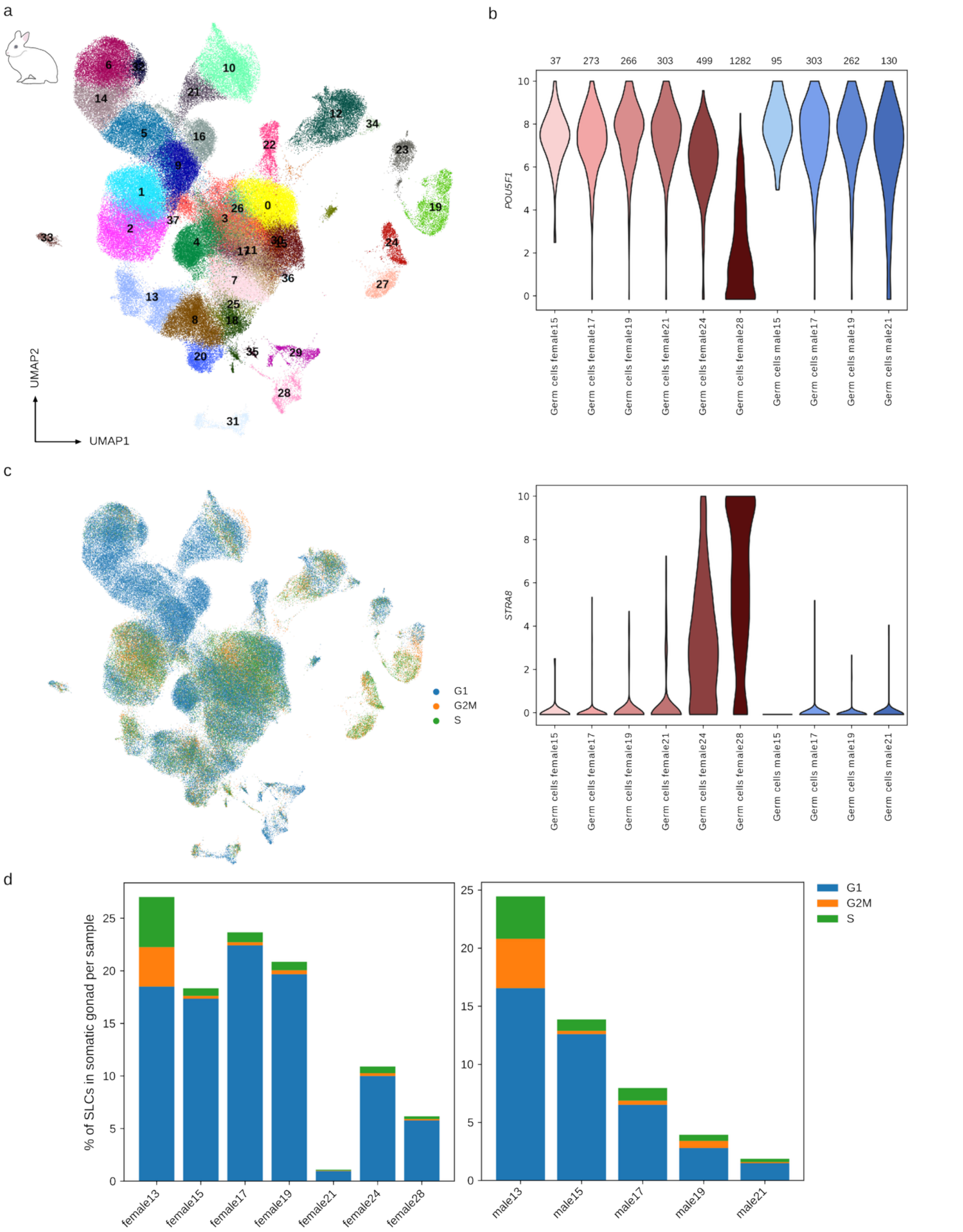


**Supplementary Figure 1. Leiden clustering, cell cycle scores and germ cells characterization.** a) UMAP projection of the integrated samples coloured by Leiden clusters. b) Violin plots displaying the expression of *POU5F1* (upper plot) and *STRA8* (lower plot) in germ cells of different stages. Stages with less than 20 germ cells are not shown. The number of germ cells in each stage is indicated in the upper part of the plot. c) UMAP projection coloured by cell cycle phase. d) Barplot displaying the percentage of SLCs at different cell cycle phases in the somatic gonad (mesonephros and other extragonadal cell types not included) in the female (left plot) and in the male (right plot).

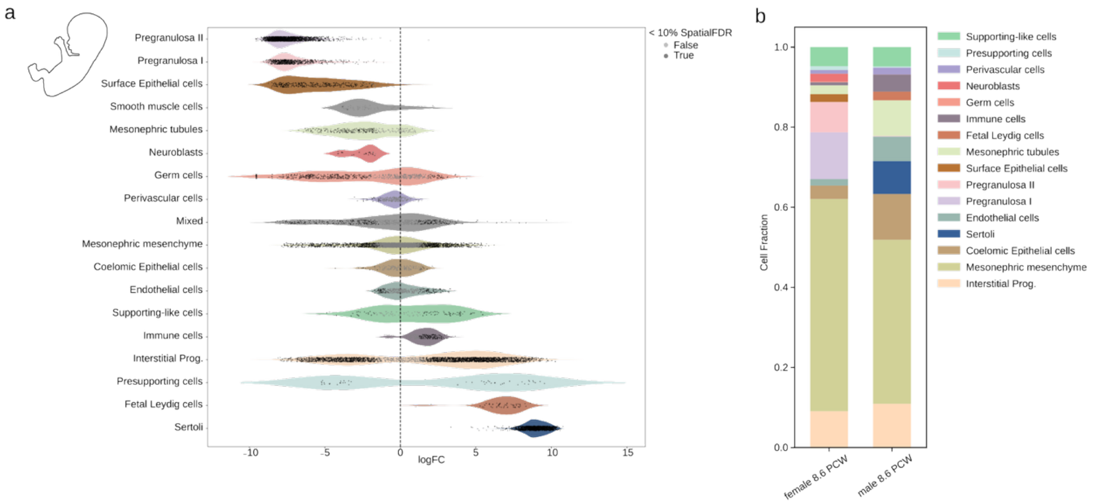


**Supplementary Figure 2. Differential abundance analysis of human gonadal cell types**. a) Violin plot showing the logFC of the milo neighbourhoods per cell type in human data. A negative logFC means that the neighbourhood is enriched in female cells and a positive logFC in male cells. Neighbourhoods displaying significant differential abundance between sexes (spatialFDR < 0.1) are shown in dark grey, whereas not significant ones are represented in light grey. A mild male bias was observed for the human SLCs. b) Barplot showing the human cell type proportions at 8.6 PCW, approximately equivalent to mouse 13.5 d*pc*.

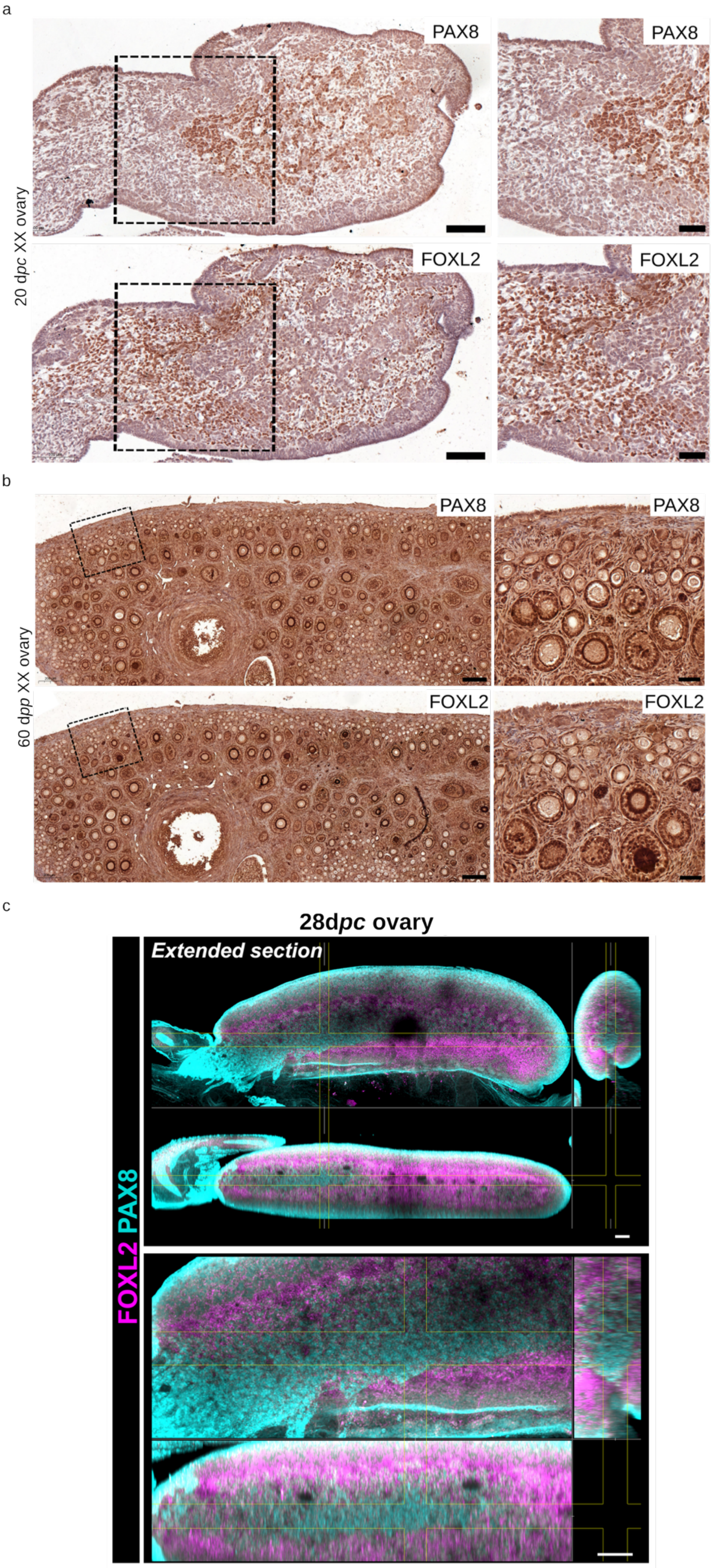

a
20 dpc XX ovary
PAX8
PAX8
FOXL2
FOXL2
b
60 dpp XX ovary
PAX8
PAX8
FOXL2
FOXL2
c
28dpc ovary
Extended section
FOXL2 PAX8

**Supplementary Figure 3. PAX8 and FOXL2 expression in control ovaries.** (a) Immunostaining of PAX8 and FOXL2 on XX control ovaries at 20 d*pc*. The dotted square indicates the enlarged area on the right. Scale bars = 100 µm on the left; 50 µm on the right. (b) Immunostaining of PAX8 and FOXL2 on XX control ovaries at 60 days *post-partum* (d*pp*). The dotted square indicates the enlarged area on the right. Scale bars = 200 µm on the left; 50 µm on the right. c) *In toto* immunofluorescence images of ECi-cleared prenatal rabbit ovary at 28 d*pc*. Bottom panel is a magnified view of the top panel. For each panel, the top image shows an extended projection of the XY plane, the bottom image shows an extended projection of the XZ plane, and the right image shows an extended projection of the ZY plane. PAX8 is labeled in cyan and FOXL2 in magenta. Scale bars, 100 µm.

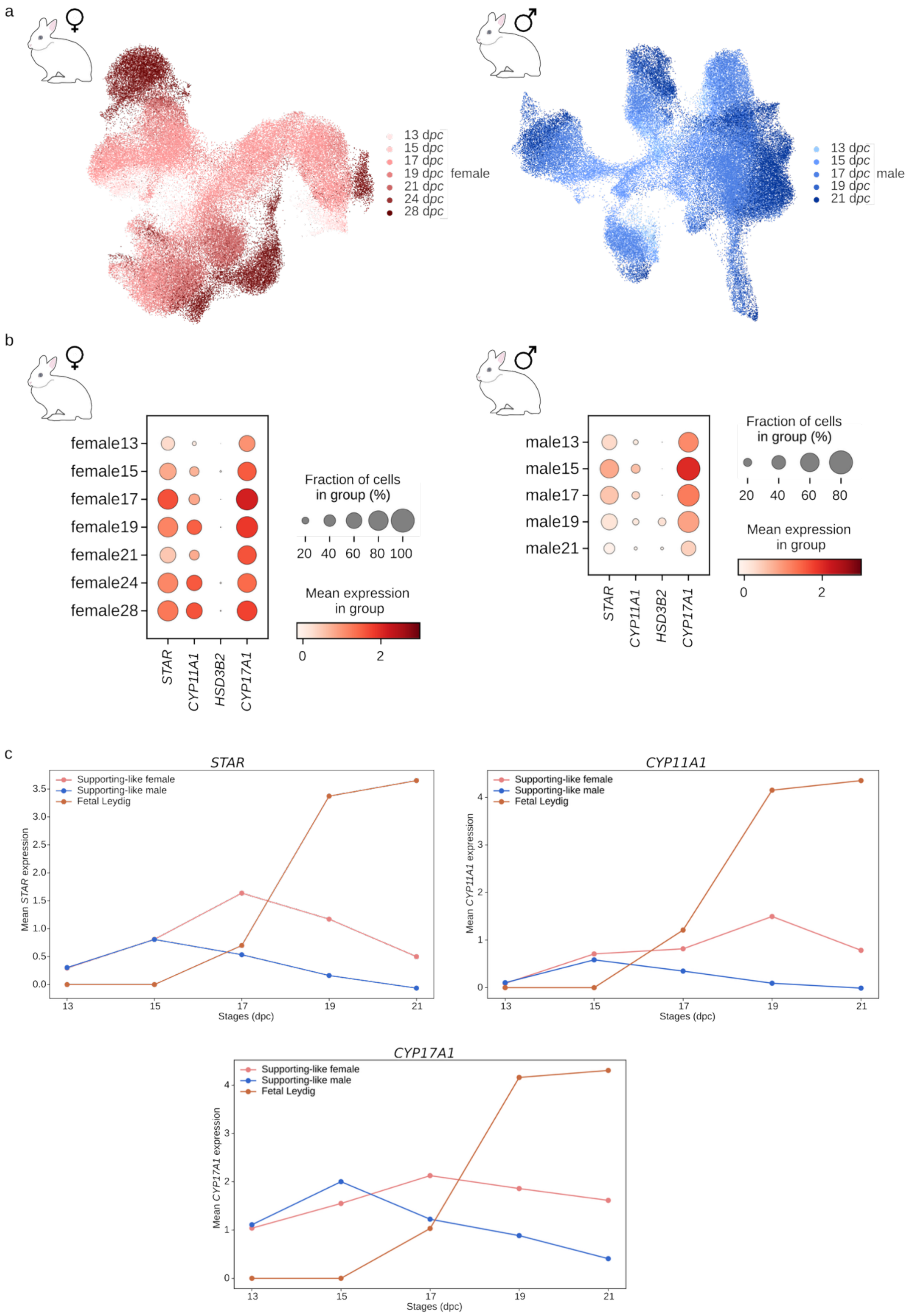


**Supplementary Figure 4. Expression of the androgen synthesis genes.** a) UMAP projection colored by female (left plot) and male (right plot) stages. b) Dotplot displaying the mean scaled expression of *STAR*, *CYP11A1*, *HSD3B2* and *CYP17A1* across stages in the SLCs of female (left plot) and male (right plot). c) Mean scaled expression of *STAR* (left plot), *CYP11A1* (right plot) and *CYP17A1* (bottom plot) in female and male SLCs and in fetal Leydig cells across stages.

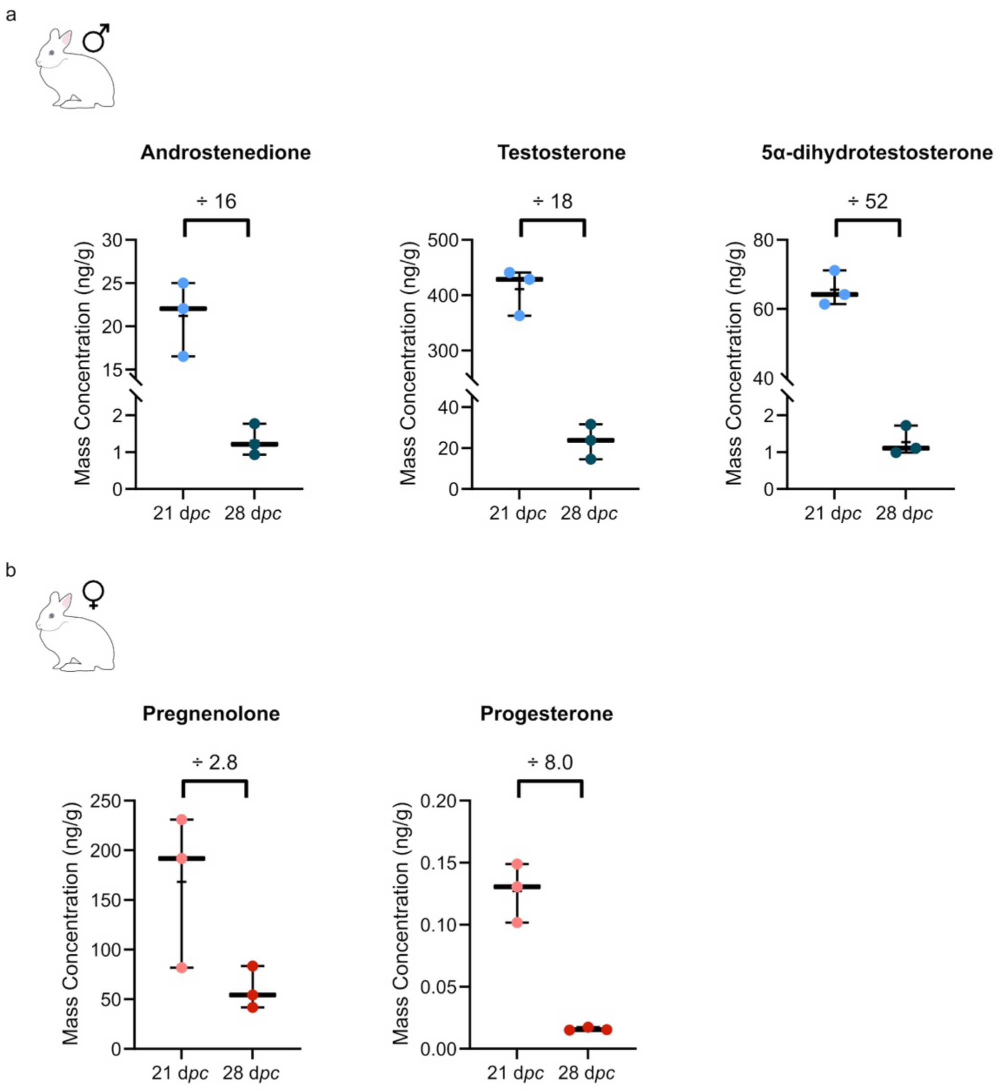


**Supplementary Figure 5**. Steroids concentrations in female and male rabbit gonads. a) Androstenedione, testosterone and 5α-dihydrotestosterone concentrations in male rabbit testes. b) Pregnenolone and progesterone concentrations in female rabbit ovaries. Two-tailed t-tests were performed for evaluating significance.

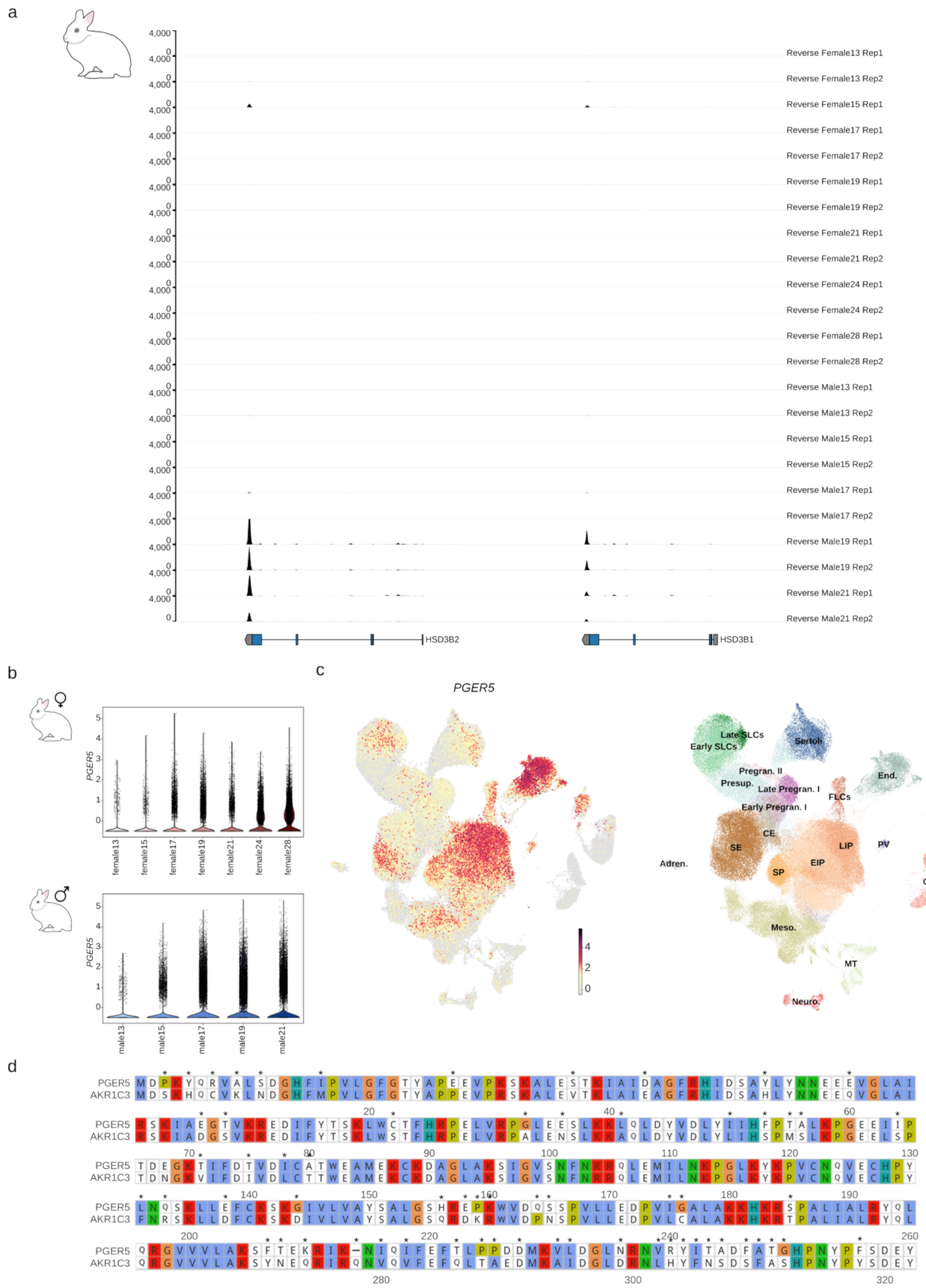


**Supplementary Figure 6.** Expression of *HSD3B2*, *HSD3B1* and *PGER5* across samples and alignment of rabbit PGER5 protein with human AKR1C3 protein. a) Expression tracks of the reverse strand for all the libraries for the region chr13:45107813-45159285 encompassing the *HSD3B2* and *HSD3B1* genes. High expression is observed in the male libraries at 19 and 21 d*pc*. No expression is observed in the female samples with the exception of some residual expression at 15 d*pc* coming from adrenal gland cells. b) Violin plots displaying the expression of *PGER5* in the female (upper plot) and in the male (lower plot) stages. c) UMAP projection showing the expression of *PGER5* in the full rabbit dataset. d) Protein alignment between rabbit PGER5 and human AKR1C3. Sequence mismatches are highlighted with asterisks. Early SLCs, early supporting-like cells; Late SLCs, late supporting-like cells; CE, coelomic epithelial cells; SE, surface epithelial cells; Presup., presupporting cells; Sertoli, Sertoli cells; Pregran. II, pregranulosa II cells; Early Pregran. I, early pregranulosa I cells; Late Pregran. I, late pregranulosa I cells; EIP, early interstitial/stromal progenitors; LIP, late interstitial progenitors; SP, stromal progenitors; FLC, fetal Leydig cells; PV, perivascular cells; Imm., immune cells; Meso., mesonephric mesenchymal cells; MT, mesonephric tubules; End., endothelial cells; GC, germ cells; EC, erythrocytes; Neuro., neuroblasts; Adren., adrenal gland cells.

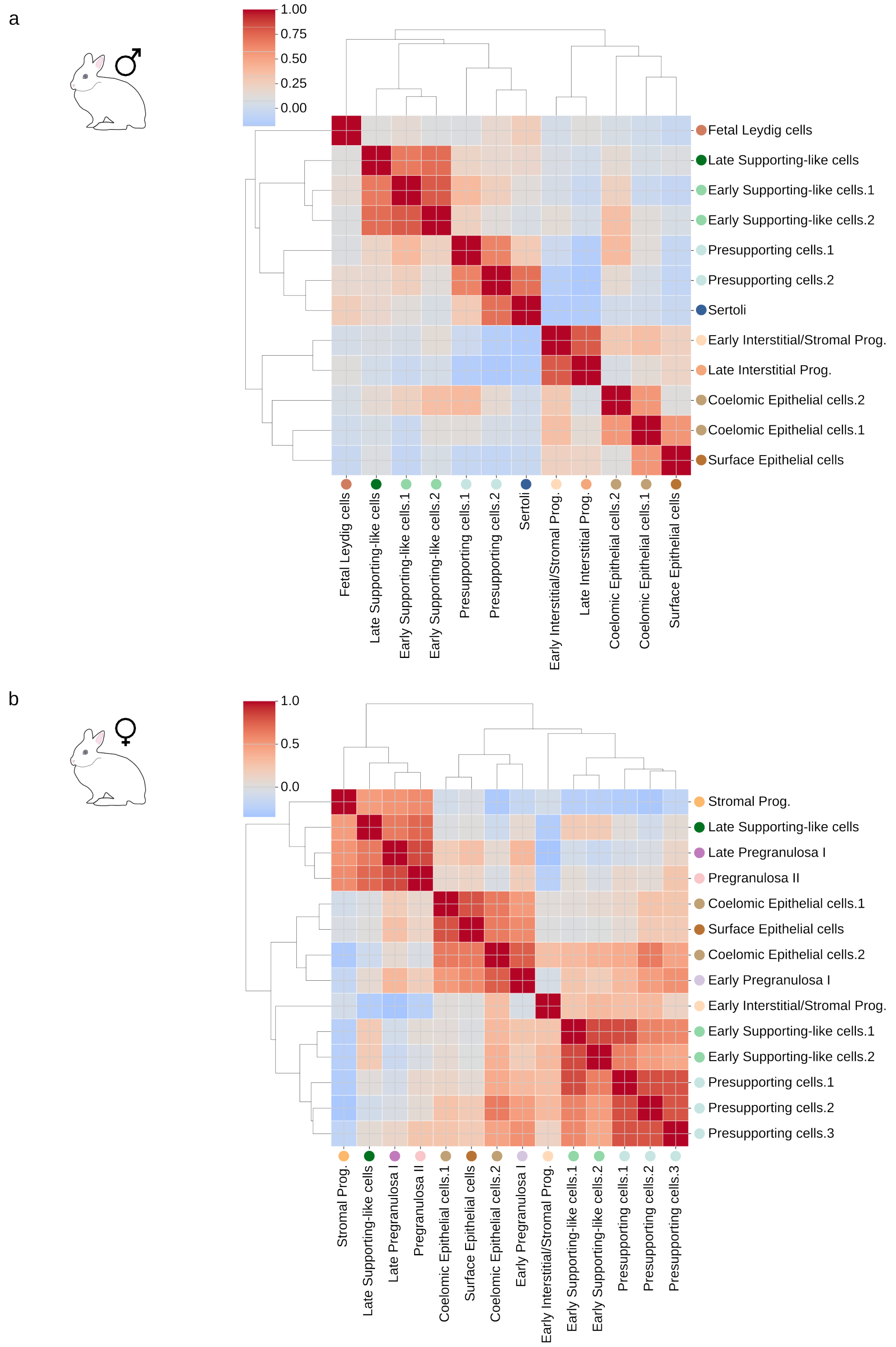


**Supplementary Figure 7**. Correlation of transcriptomes. a-b) Heatmaps displaying the correlation between transcriptomes of (a) male and (b) female cell types.

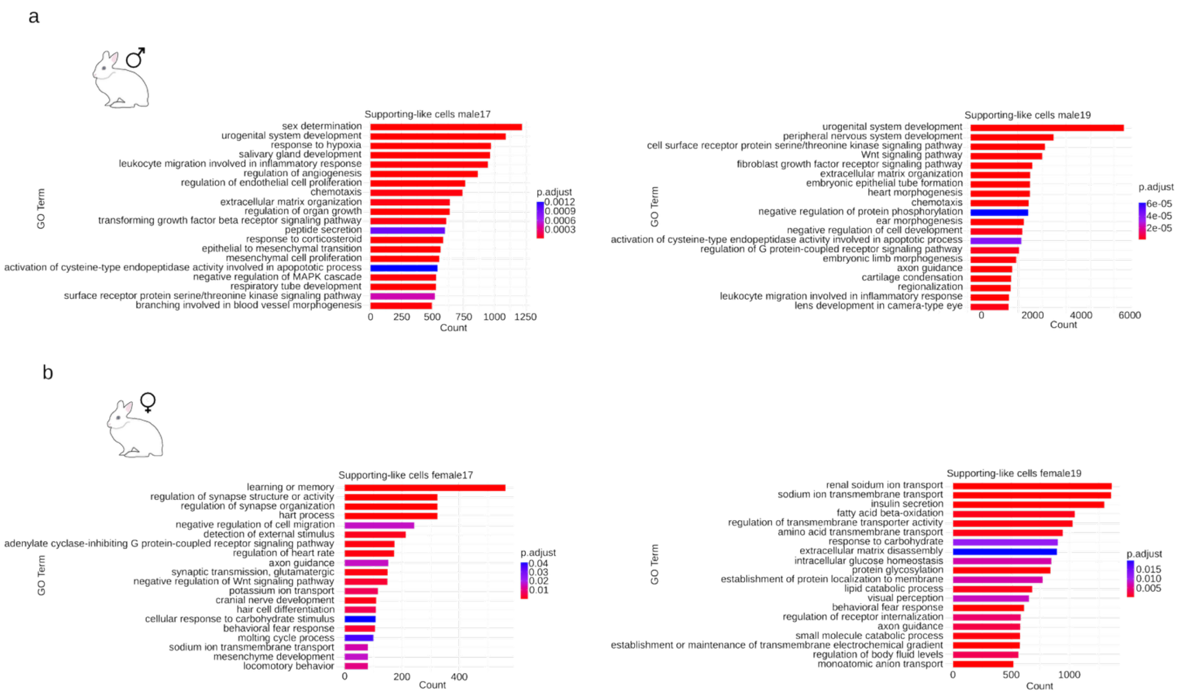


**Supplementary Figure 8**. GO enrichment analysis for female and male SLCs. a) Barplot highlighting GO terms significantly enriched in male SLCs vs female SLCs at 17 (left plot) and 19 (right plot) d*pc*. b) Same barplots that in (a) for GO terms significantly enriched in female SLCs vs male SLCs.

a

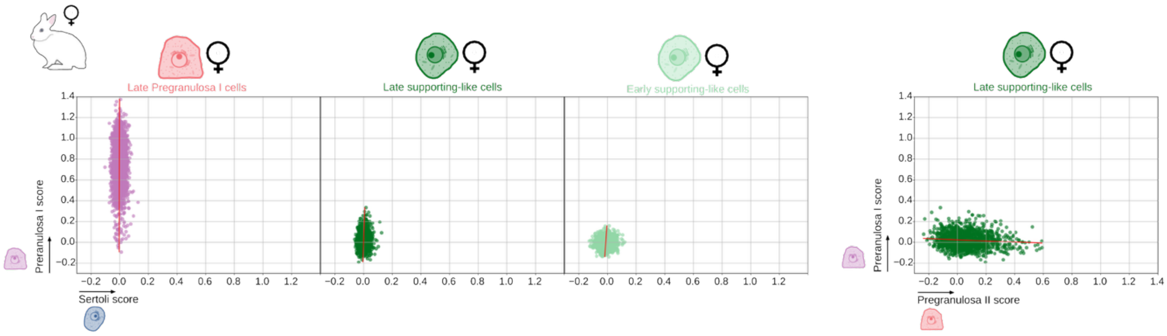


**Supplementary Figure 9**. Transcriptional characterization of Pregranulosa I and female SLCs. a) Scatterplots showing the pregranulosa I vs Sertoli scores (left plots) and pregranulosa I vs pregranulosa II scores (right plot) from LASSO models for female cell types.

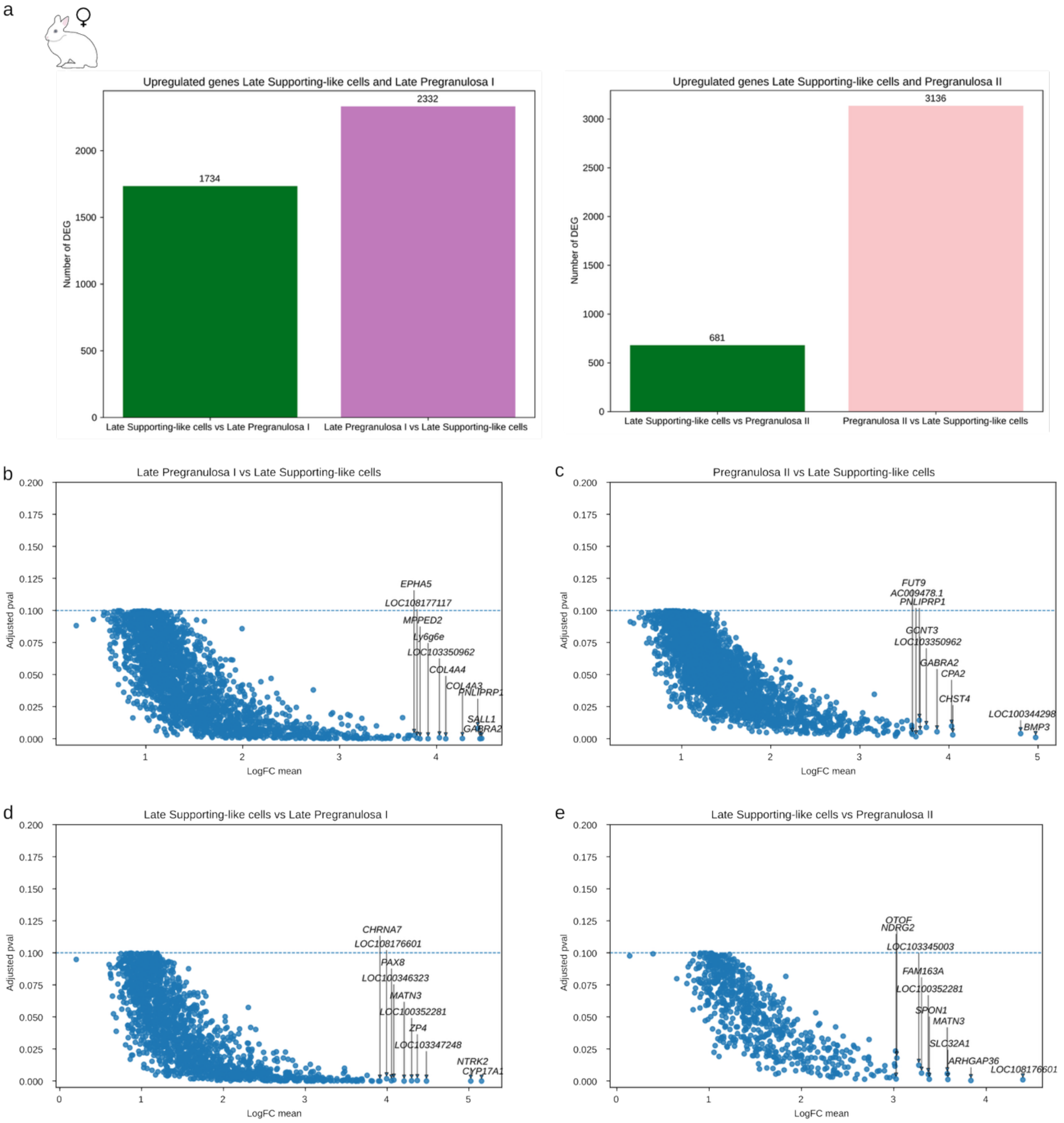


**Supplementary Figure 10**. Upregulated genes in female SLCs and Pregranulosa populations. a) Barplot showing the number of upregulated genes between late SLCs and late Pregranulosa I (left plot) and late SLCs and Pregranulosa II (right plot). b-e) Scatter plots highlighting the top 10 significant upregulated genes based on logFC in late Pregranulosa I for the late Pregranulosa I vs late SLCs comparison (b), top 10 genes in Pregranulosa II for the Pregranulosa II vs late SLCs comparison (c), top 10 genes in late SLCs for the late SLCs vs late Pregranulosa I comparison (c) and top 10 genes in late SLCs for the late SLCs vs Pregranulosa II comparison (d). An adjusted p-value threshold of 0.1 was established.

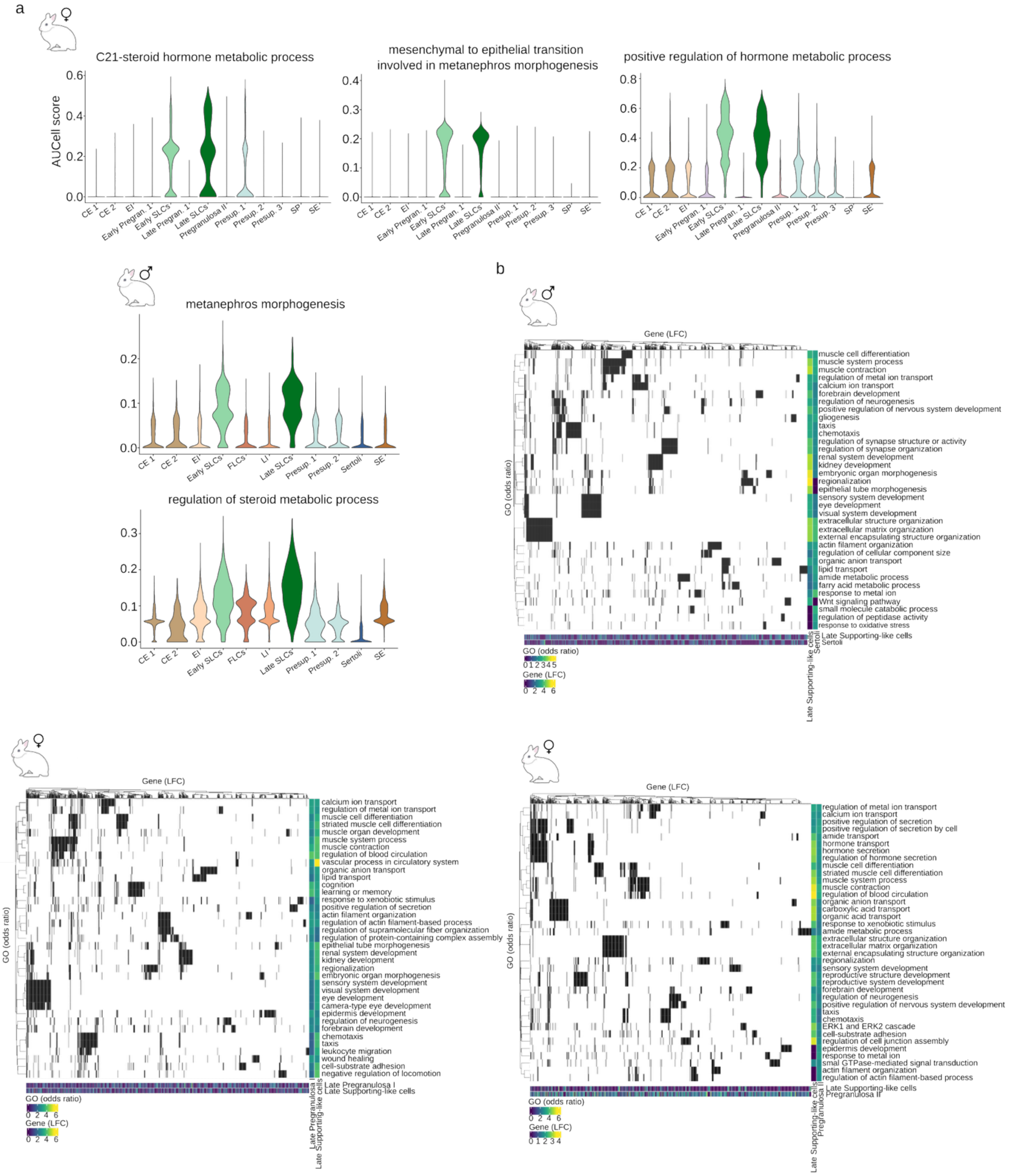


**Supplementary Figure 11**. GO enrichment analysis results for SLCs, Pregranulosa and Sertoli. a) AUCell scores for the genes included in different GO terms. b) Heatmaps displaying GO terms significantly enriched in either late SLCs, late Pregranulosa I, Pregranulosa II or Sertoli upregulated genes of the comparatives: (upper plot) Late SLCs vs Sertoli cells, (lower left plot) Late SLCs vs Late Pregranulosa I and (lower right plot) Late SLCs vs Pregranulosa II. In the Y axis the GO terms can be observed coloured by their odds ratio in both cell types. In the X axis the genes coloured by their logFC in each of the cell types. Early SLCs, early supporting-like cells; Late SLCs, late supporting-like cells; CE, coelomic epithelial cells; SE, surface epithelial cells; Presup., presupporting cells; Sertoli, Sertoli cells; Pregran. II, pregranulosa II cells; Early Pregran. I, early pregranulosa I cells; Late Pregran. I, Late pregranulosa I cells; EI, early interstitial/stromal progenitors; LI, late interstitial progenitors; SP, stromal progenitors; FLC, fetal Leydig cells.

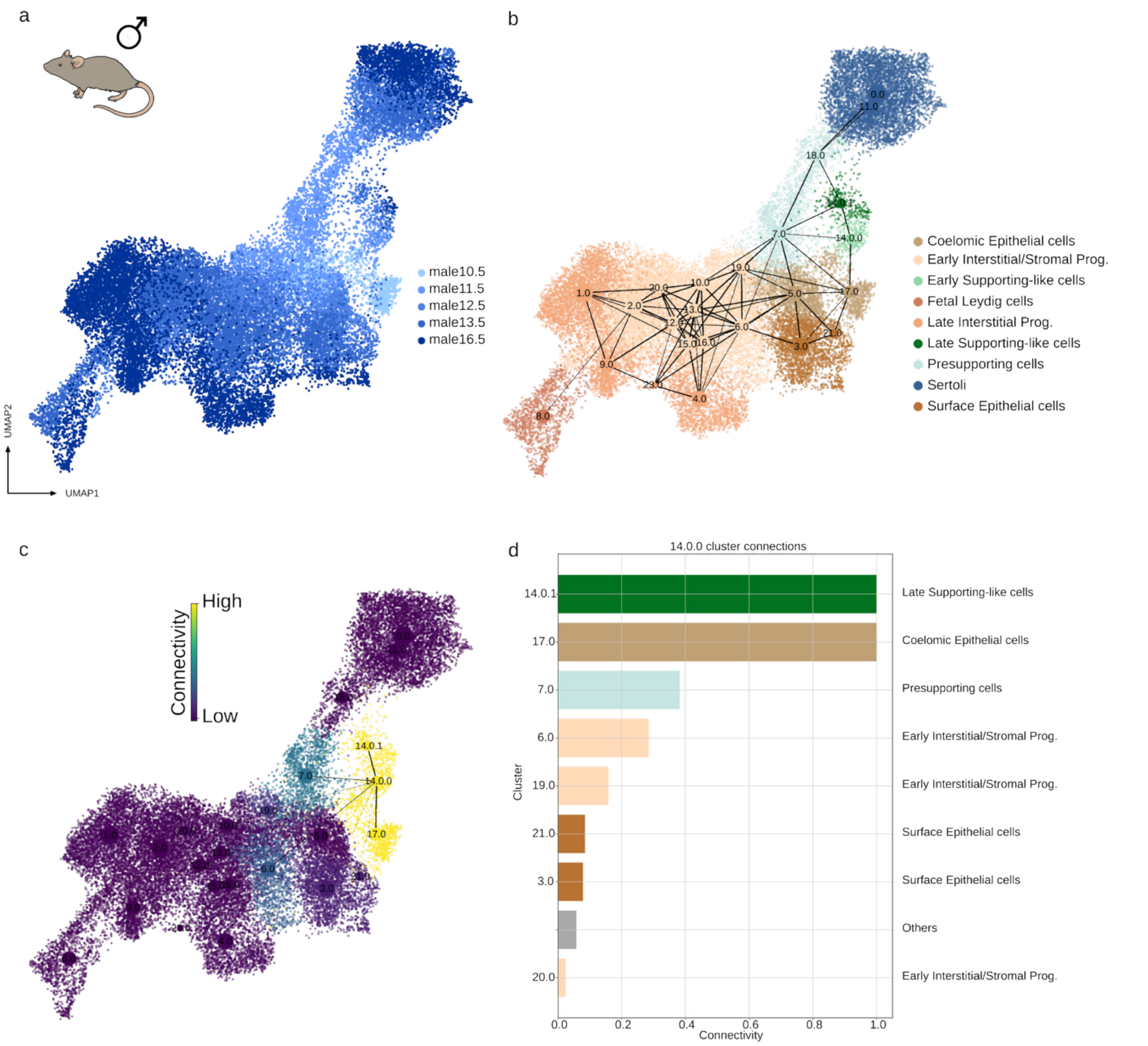


**Supplementary Figure 12**. PAGA connectivity graph of male mouse somatic gonad. a) UMAP projection of male mouse somatic gonad cells coloured by stage. b) PAGA graph over the same UMAP projection that (a). The width of the connections represents the connectivity inferred by PAGA and potentially reflects cell type similarities. c-d) PAGA connections restricted to the early SLCs cluster (cluster 14.0.0) highlighted over the UMAP projection (c). The intensity of the colour in the UMAP represents the connectivity of the clusters to the cluster 14.0.0 and it is equivalent to the length of the bars in the barplot (d).

a

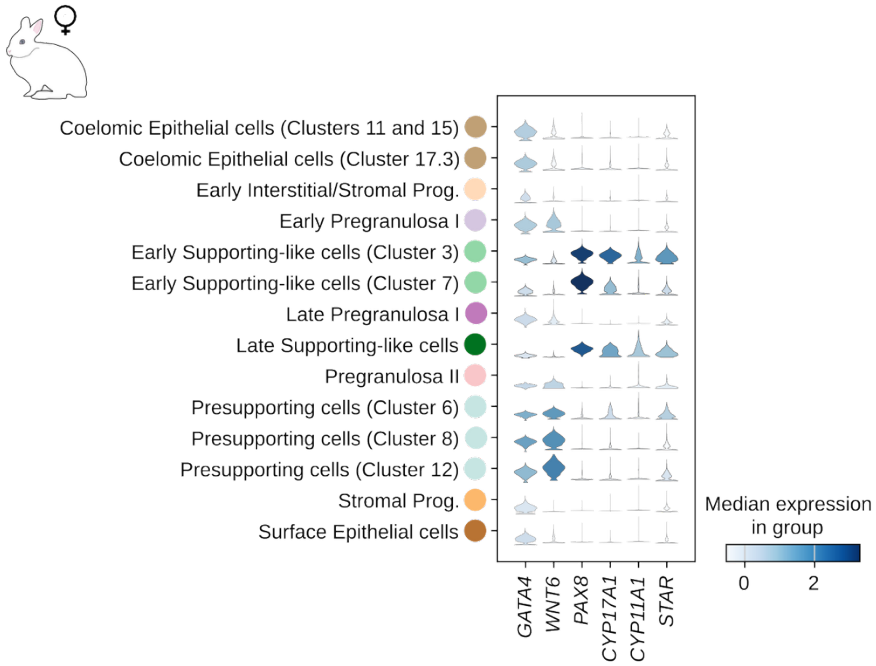


**Supplementary Figure 13**. Expression of *GATA4*, *PAX8*, *WNT6* and the androgen synthesis genes in female rabbit somatic gonad. a) Stacked violin displaying the expression of *GATA4*, *WNT6*, *PAX8* and the androgen synthesis genes for the cell types present in the somatic gonad of female rabbits.

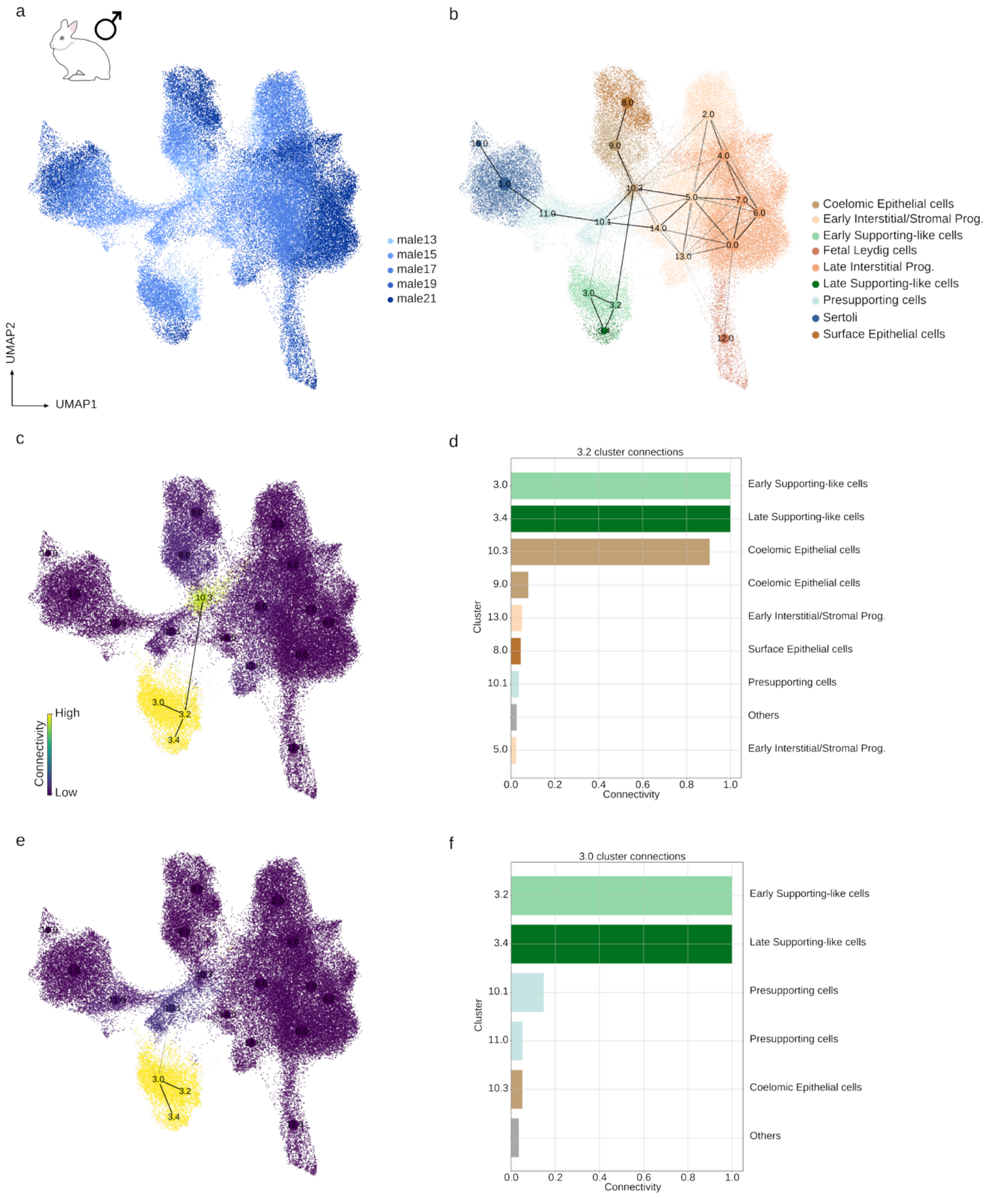


**Supplementary Figure 14**. PAGA connectivity graph of male rabbit somatic gonad. a) UMAP projection of male rabbit somatic gonad cells coloured by stage. b) PAGA graph over the same UMAP projection that (a). The width of the connections represents the connectivity inferred by PAGA and potentially reflects cell type similarities. c-d) PAGA connections restricted to one of the early SLCs cluster (cluster 3.2) highlighted over the UMAP projection (c). The intensity of the colour in the UMAP represents the connectivity of the clusters to the cluster 3.2 and it is equivalent to the length of the bars in the barplot (d). e-f) Same that (c-d) for the other early SLC cluster (cluster 3.0).

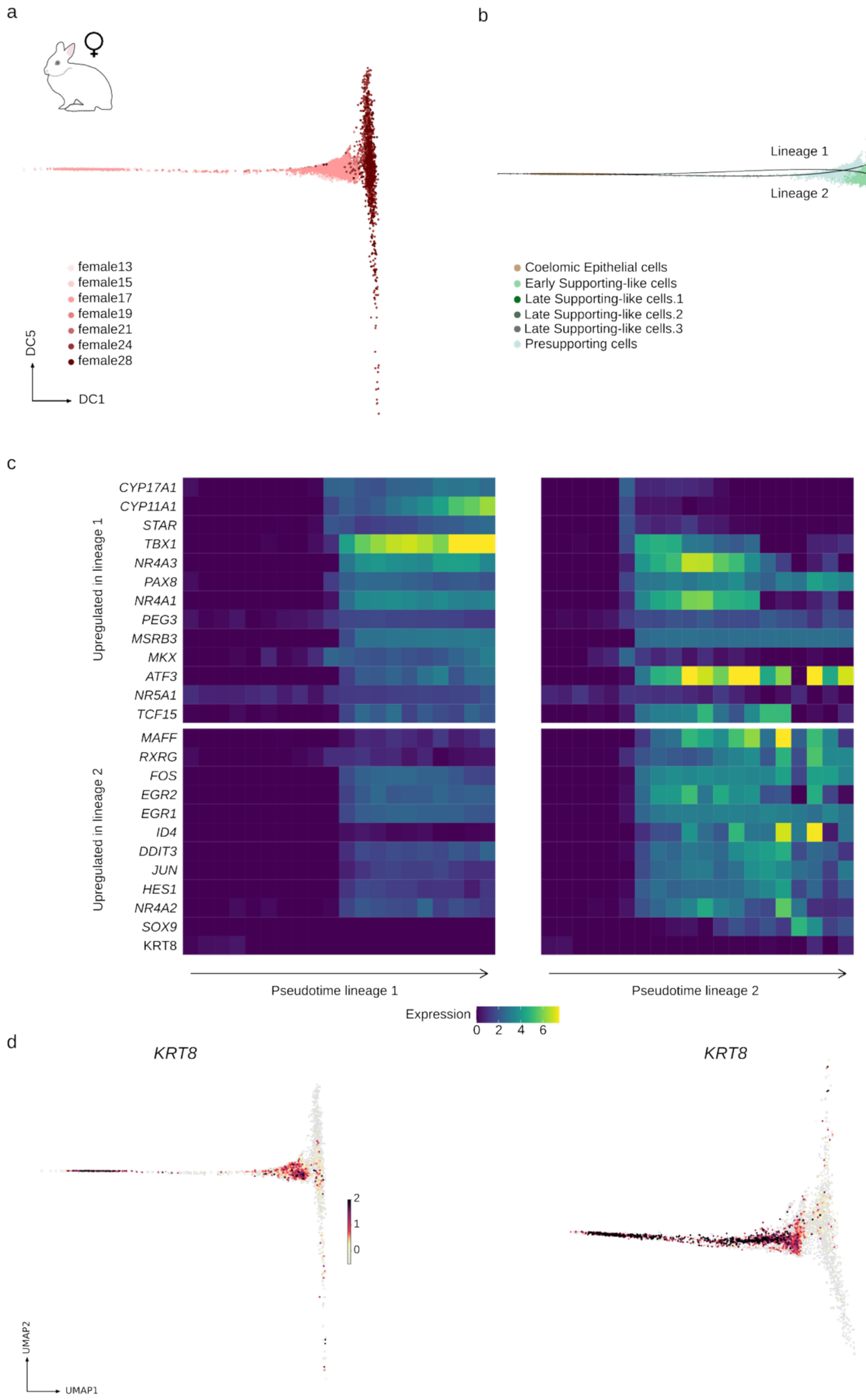


**Supplementary Figure 15**. Female SLCs trajectory from presupporting cells. a-b) Diffusion maps of the PAGA highly connected clusters starting from the coelomic epithelium (cluster 15), passing through the presupporting cells (clusters 6 and 8) and ending in late SLCs for the female rabbit, coloured by stages (a) and by cell type (b). The same subpopulations of late SLCs as in the coelomic epithelium trajectory were identified. Slingshot was used to order the cells along two lineages and to compute a pseudotime. c) Heatmap showing gene expression trends along the slingshot lineages for the androgen-synthesis genes, upregulated genes in at least one of the lineages (from the coelomic epithelium trajectory as in Figure 6), *SOX9* and

*KRT8*. Cells were separated in 20 blocks based on their pseudotime value. The colours represent the mean expression value in each block. Equivalent tendencies to those of the trajectory from the coelomic epithelium can be observed. d) Diffusion maps of the presupporting trajectory (left plot) and the coelomic epithelium trajectory (right plot) coloured by the expression of *KRT8*. To facilitate visualization, cells with no detectable expression are shown in grey.

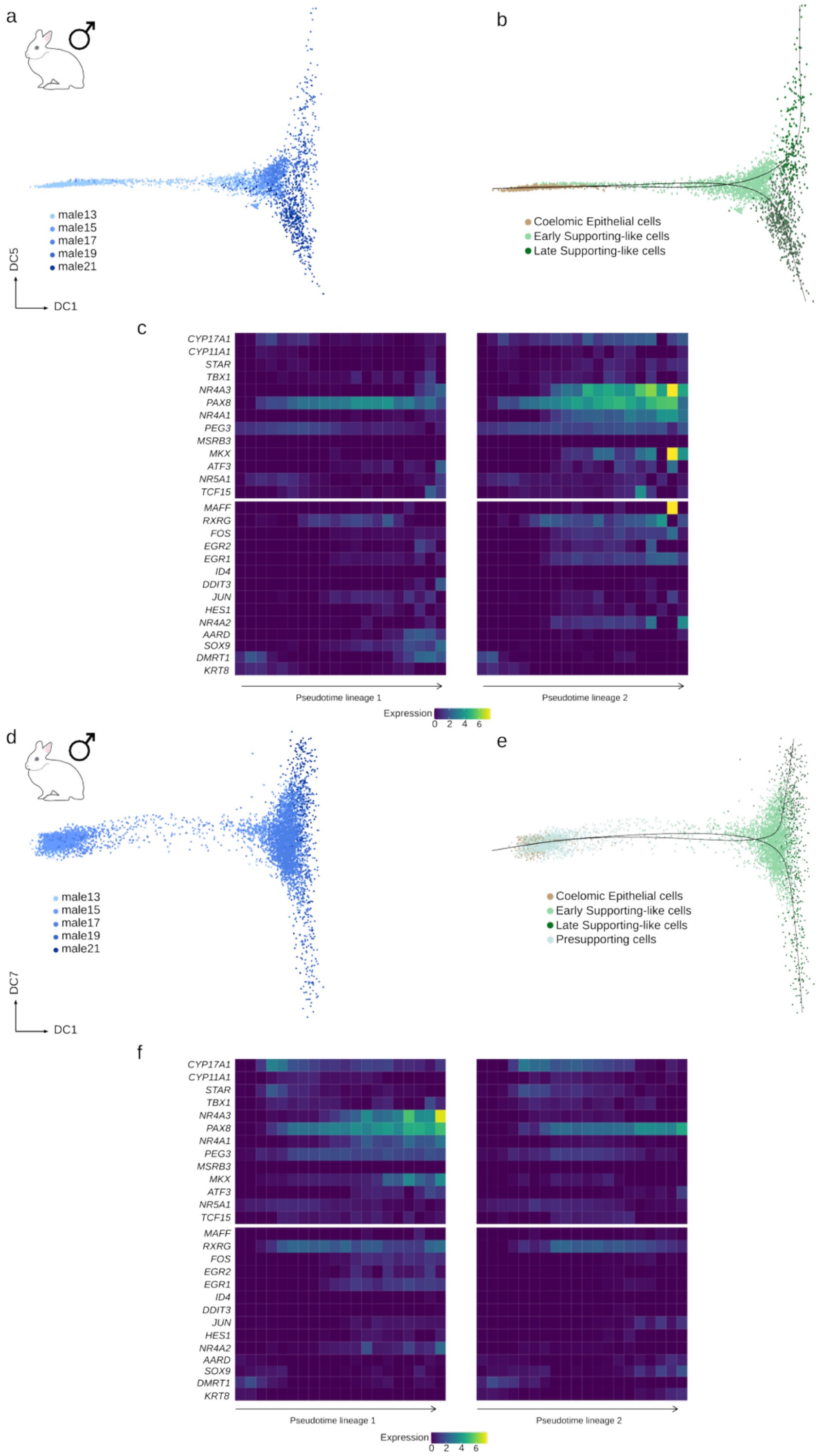


**Supplementary Figure 16**. Male SLCs trajectories. a-b) Diffusion maps of the PAGA highly connected clusters starting from the coelomic epithelium (cluster 10.3) and ending in late SLCs for the male rabbit, coloured by stages (a) and by cell type (b). Slingshot was used to order the cells along two lineages and to compute a pseudotime. c) Heatmap showing gene expression trends along the slingshot lineages for the androgen-synthesis genes, upregulated genes in at least one of the lineages (from the coelomic epithelium trajectory as in Figure 4), *SOX9*, AARD, DMRT1 and *KRT8*. Cells were separated in 20 blocks based on their pseudotime value. The colours represent the mean expression value in each block. Some of the genes that are upregulated in the female trajectory are also upregulated in the male. Slightly higher expression of *CYP17A1* is observed in the lineage 2 and the opposite happens for *SOX9, AARD and DMRT1*. d-f) Same as in a-c). In (a and b) the diffusion map starts from the coelomic epithelium (cluster 10.3) passing through the presupporting cells (cluster 10.1) and ending in late SLCs for the male rabbit. In (f) the gene trends are less obvious than in (c) due to the worse resolution of the reduced dimensional space.

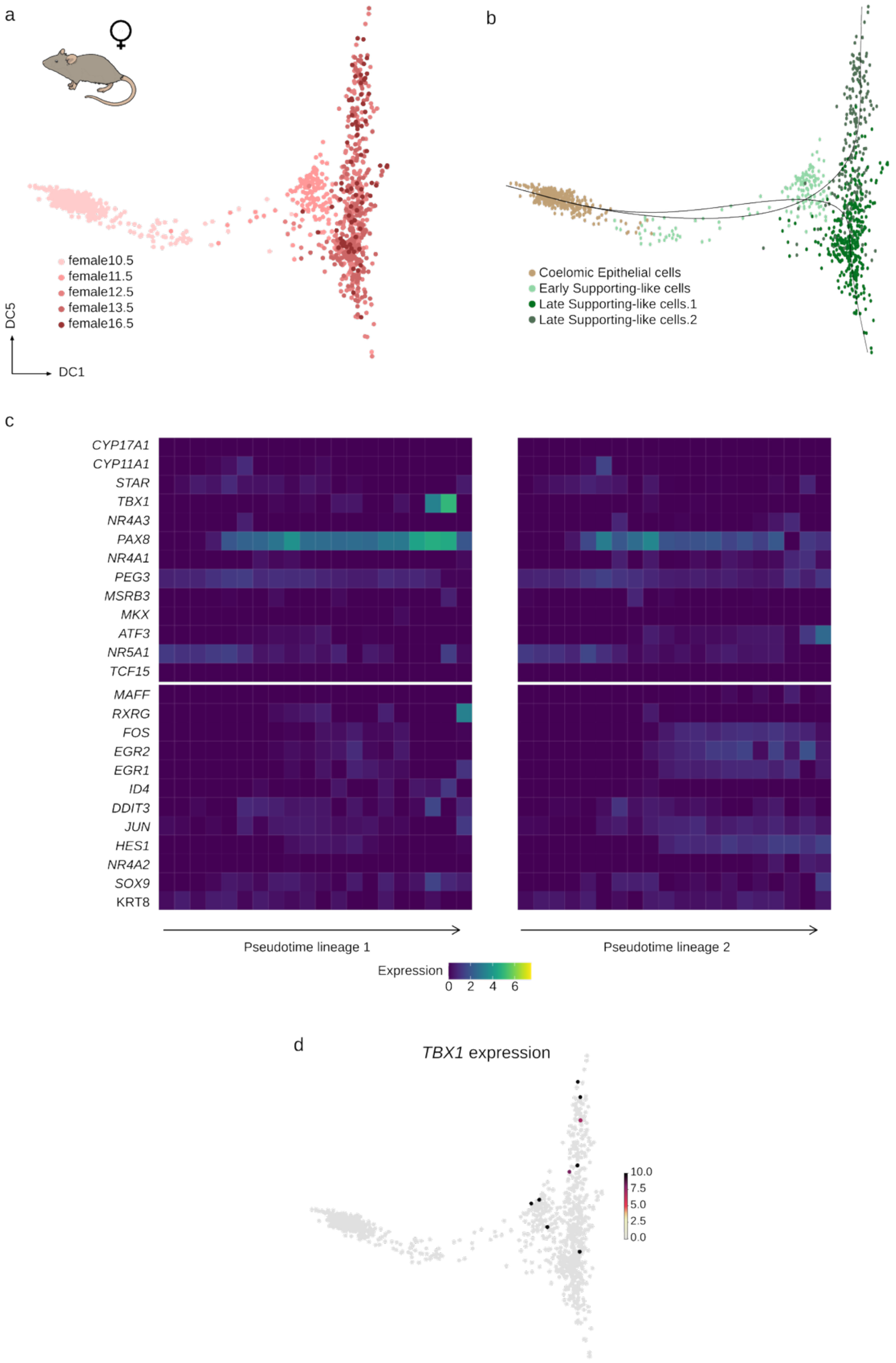


**Supplementary Figure 17**. Female mouse SLCs trajectory from coelomic epithelium cells. a-b) Diffusion maps of the PAGA highly connected clusters starting from the coelomic epithelium (cluster 7) and ending in late SLCs for the female mouse, coloured by stages (a) and by cell type (b). Slingshot was used to order the cells along two lineages and to compute a pseudotime. c) Heatmap showing gene expression trends along the slingshot lineages for the androgen-synthesis genes, upregulated genes in at least one of the lineages (from the rabbit coelomic epithelium trajectory as in Figure 6), *SOX9* and *KRT8*. Cells were separated in 20 blocks based on their pseudotime value. The colours represent the mean expression value in each block. In contrast to the rabbit, no interesting pattern of expression was observed. Slight upregulation of *TBX1* is observed in the trajectory but, as shown in d) it is due to a very low number of cells and the data being scaled.

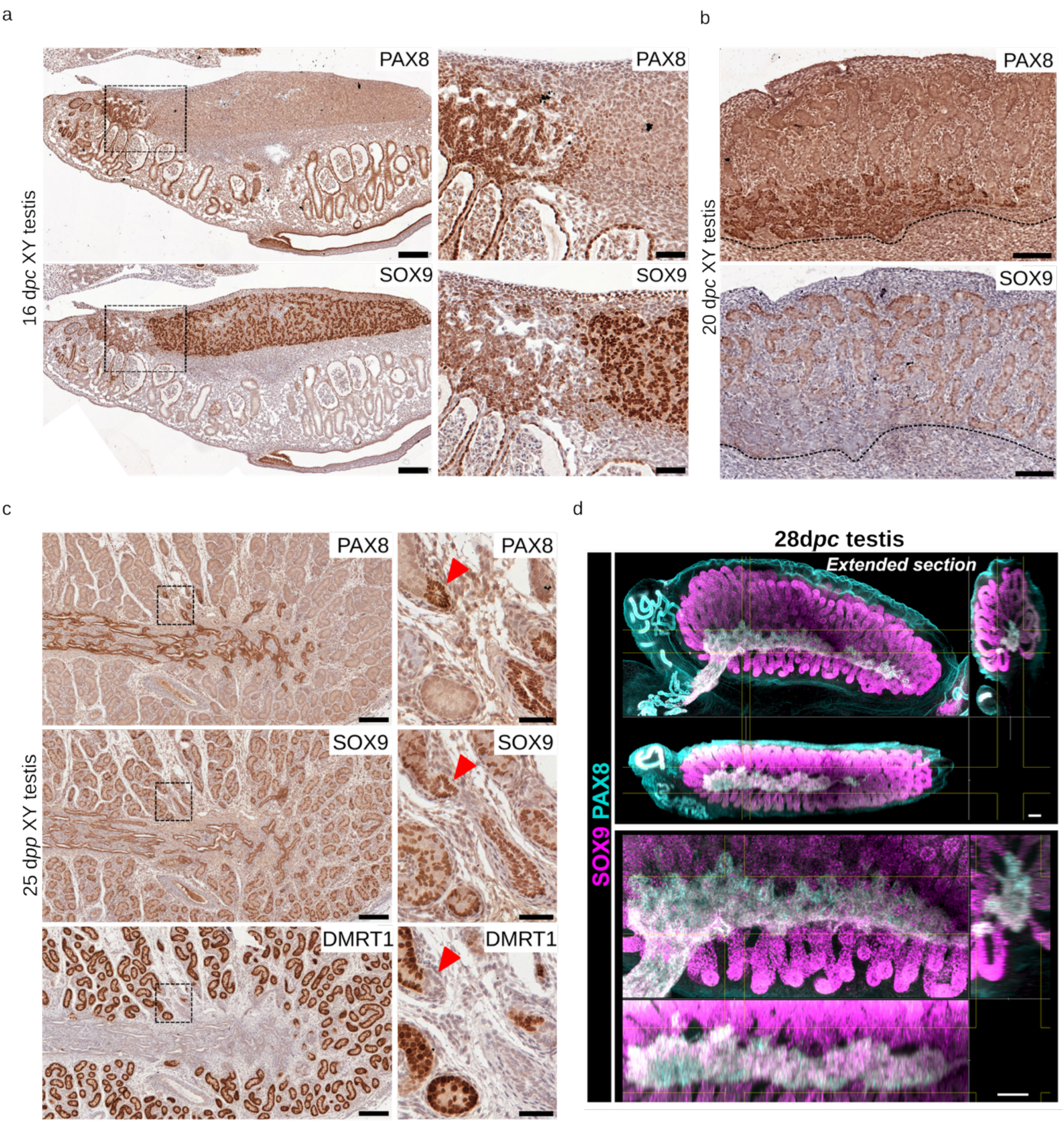


**Supplementary Figure 18**. PAX8, SOX9, and DMRT1 localization in control testes. a) Immunostaining of PAX8 and SOX9 on XY control testes at 16 d*pc*. The dotted square indicates the enlarged area on the right. Scale bars = 200 µm on the left; 50 µm on the right. b) Immunostaining of PAX8 and SOX9 on XY control testes at 20 d*pc*. The dotted line indicates the limit between the mesonephros and the gonad. Scale bars = 100 µm. c) Immunostaining of PAX8, SOX9, and DMRT1 on XY control testes at 25 d*pp*. The dotted square indicates the enlarged area on the right. Scale bars = 200 µm on the left; 50 µm on the right. The red arrowhead indicates cells positive for PAX8 and SOX9 but not for DMRT1. d) *In toto* immunofluorescence images of ECi-cleared prenatal rabbit testis at 28 d*pc*. Bottom panel is a blow-up view of the top panel. For each panel, the top image shows an extended projection of the XY plane, the bottom image shows an extended projection of the XZ plane, and the right image shows an extended projection of the ZY plane. PAX8 is labeled in cyan and SOX9 in magenta. Scale bars, 100 µm.

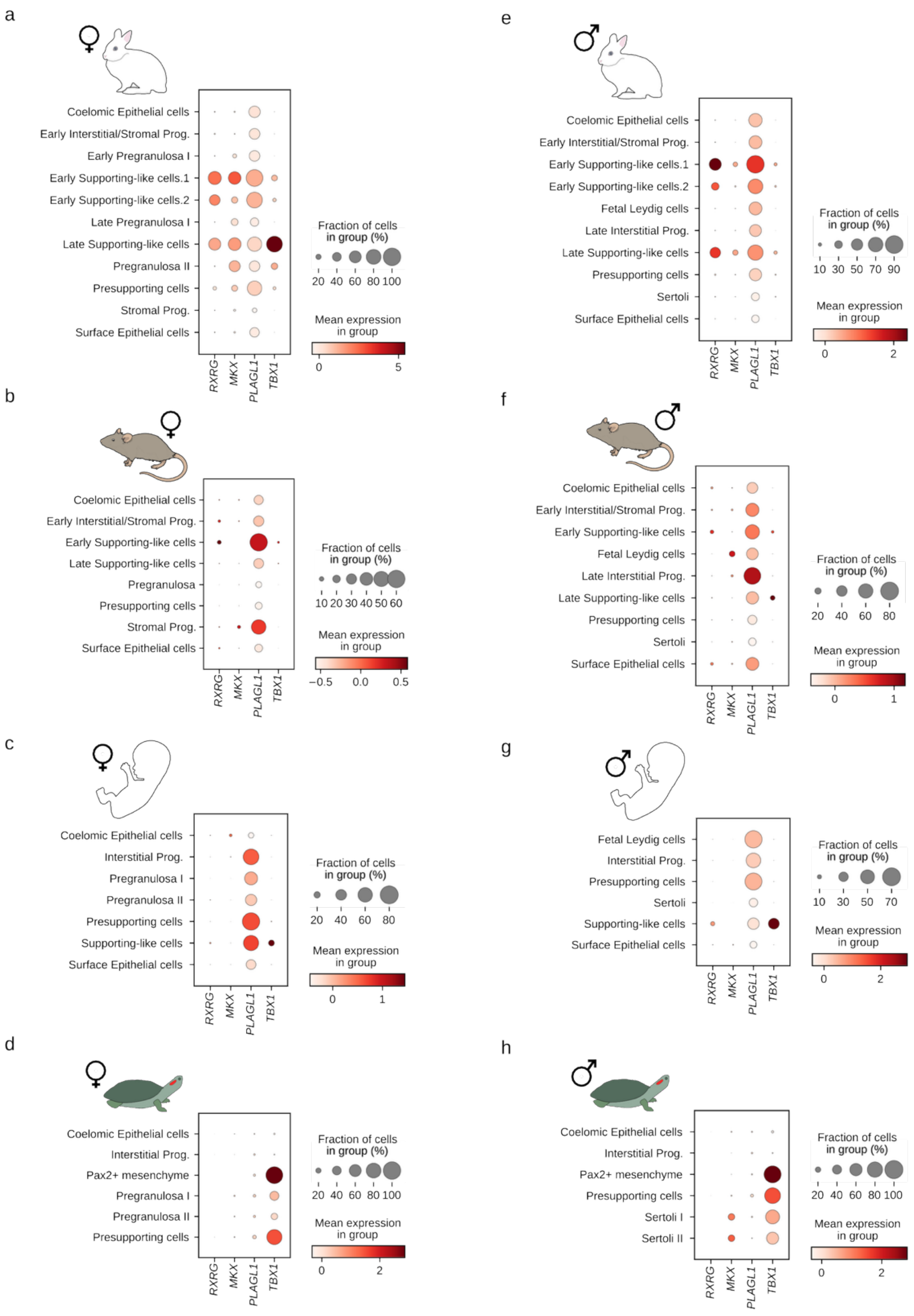


**Supplementary Figure 19**. TFs expression across species. a-d) Expression in the female datasets of rabbit (a), mouse (b), human (c) and turtle (d). e-h) Expression in the male datasets of rabbit (e), mouse (f), human (g) and turtle (h). The dot size represents the fraction of cells in the cell type expressing the gene and the intensity of the color the mean scaled expression in the cell type.

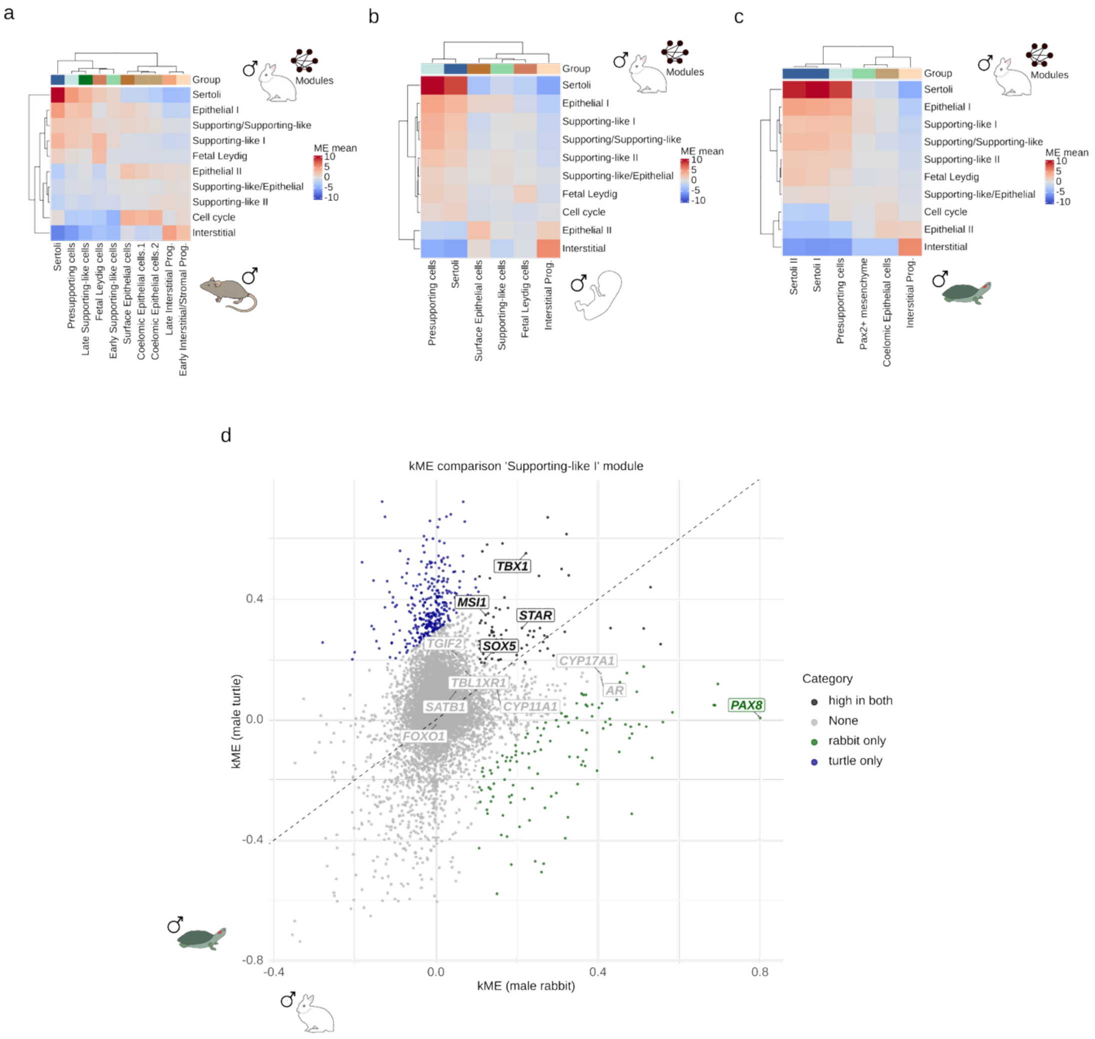


**Supplementary Figure 20**. Activity of male rabbit SLCs network across species. a-c) Heatmaps showing the ME scores of the male rabbit WGCNA modules in mouse (a), human (b) and turtle (c) cell types. This implies that the genes belonging to this module are predominantly expressed in cell types with high MEs. d) Scatterplot displaying the kMEs of the genes belonging to the rabbit SLC module in male turtle vs male rabbit. Highlighted are the same genes as in main Figure 7. In green if they are highly scored in the rabbit, in black if they are highly scored in both species and in grey if they do not surpass the threshold.

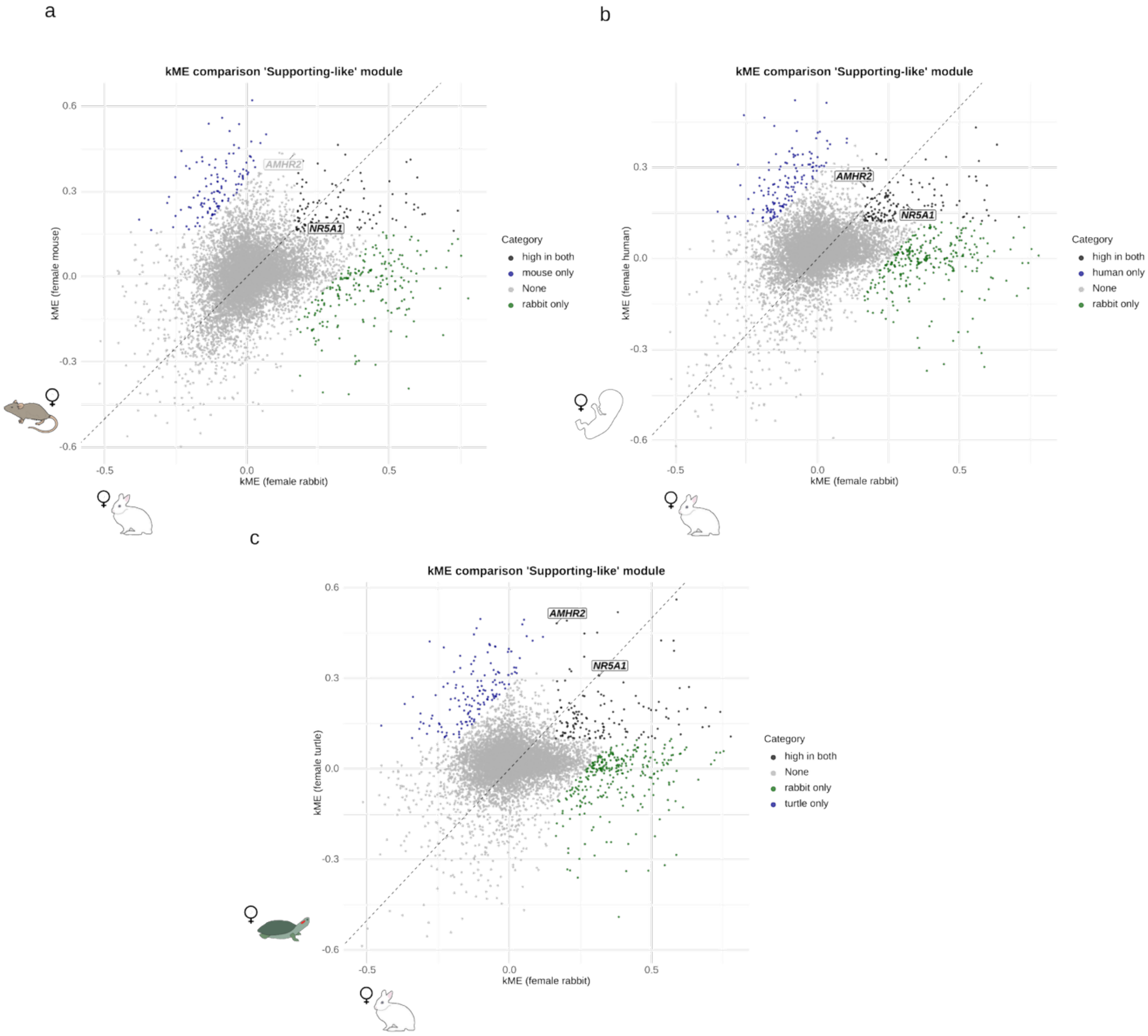


**Supplementary Figure 21**. Shared TFs of the rabbit SLCs module between species. a-c) Scatterplot displaying the kMEs of the genes belonging to the female rabbit SLC module in female mouse vs female rabbit (a), female human vs female rabbit (b) and female turtle vs female rabbit (c). Highlighted are *NR5A1* and *AMHR2*. In green if they are highly scored in the rabbit, in black if they are highly scored in both species and in grey if they do not surpass the threshold. Note that the thresholds may vary based on the dataset. Both genes surpass the threshold in all species except for *AMHR2* in the mouse comparison, in which it is close to the threshold.

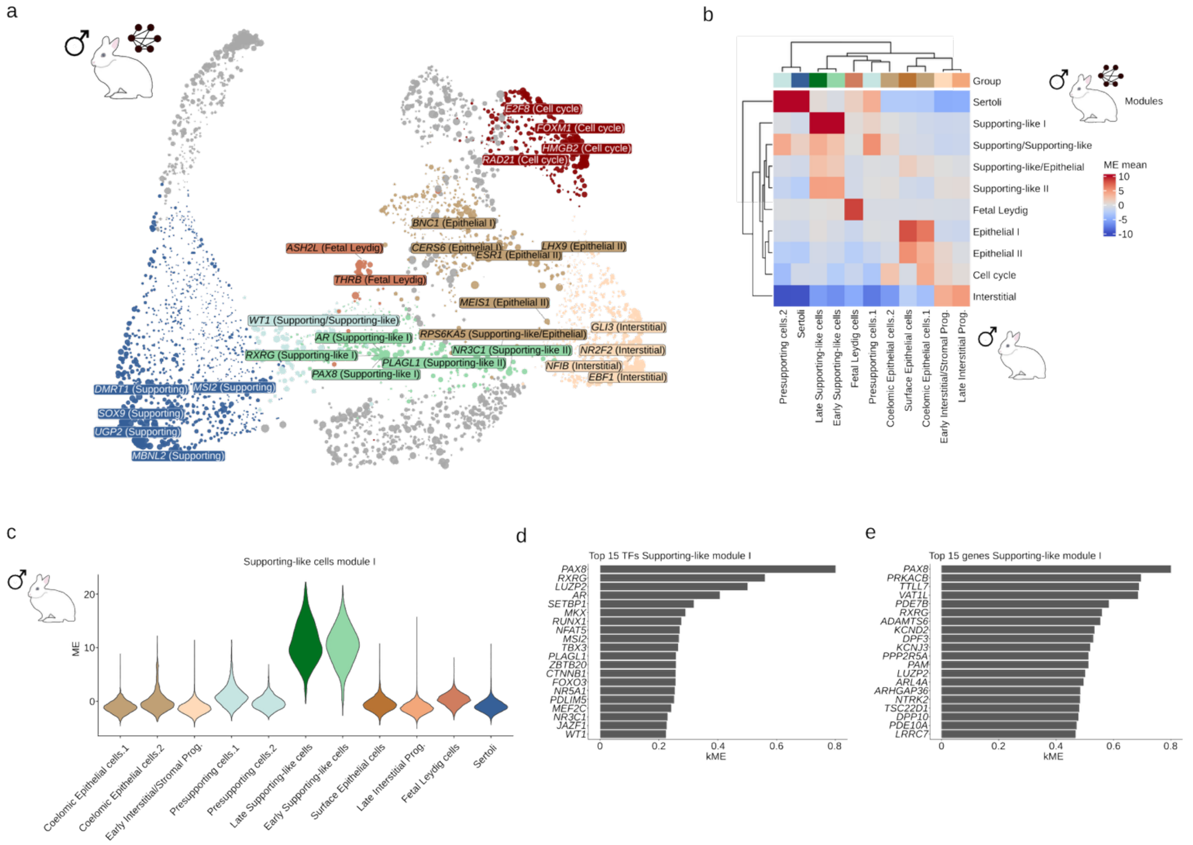


**Supplementary Figure 22**. hdWGCNA analysis on male rabbit gonads. a) UMAP projection of the male rabbit gene network computed with hdWGCNA. Genes are grouped based on their WGCNA module. The names for some top TFs per module are highlighted. b) Heatmap showing the ME scores of the male rabbit WGCNA modules in rabbit cell types.This implies that the genes belonging to this module are predominantly expressed in cell types with high MEs. c) Violin plot displaying the ME scores of the SLC module in male rabbit cell types. d-e) Barplot showing the kMEs for the top 15 TFs (d) and the top 15 genes (e) of the Supporting-like module. The higher the kME the more the expression of a gene correlates with the general expression pattern of a module.

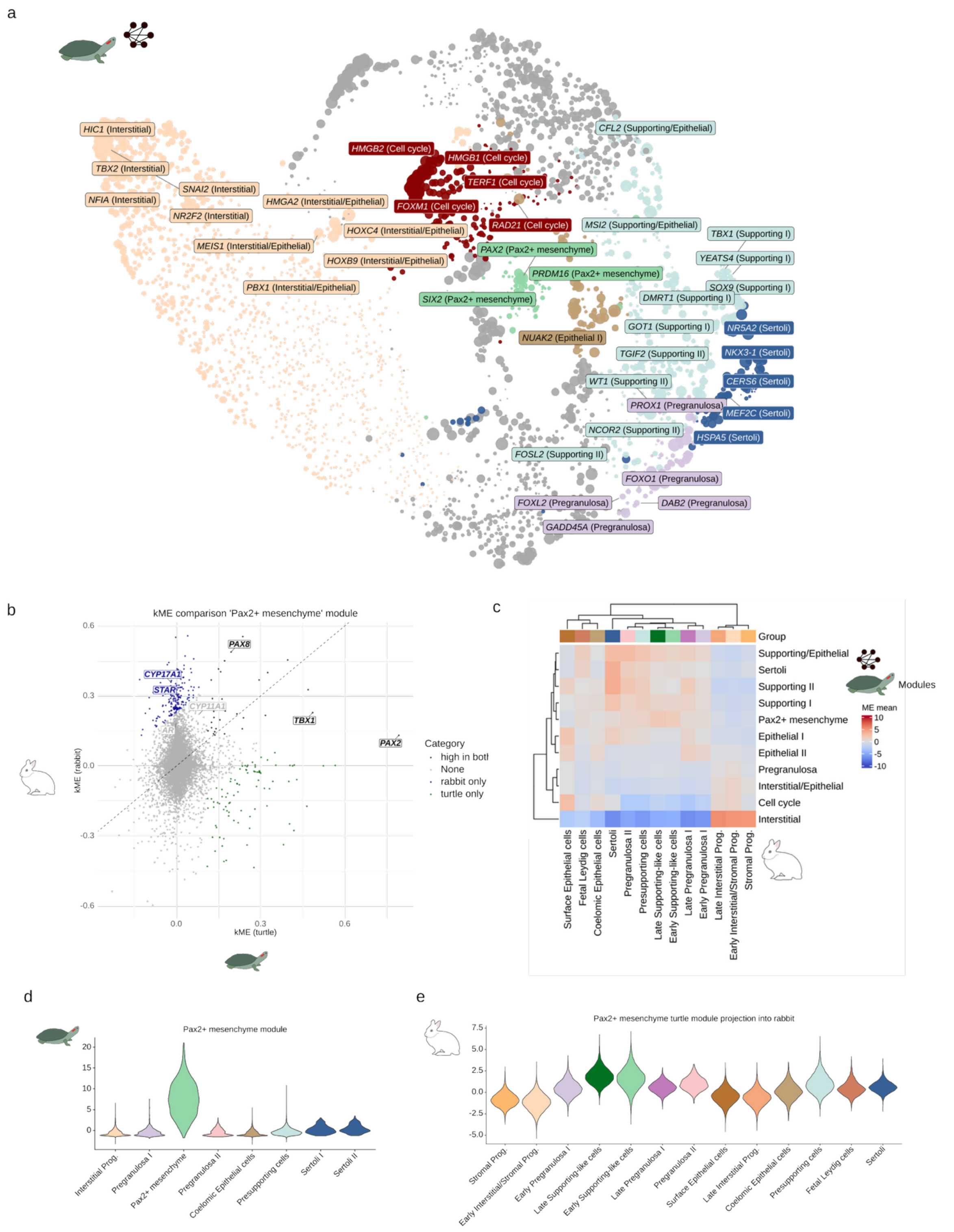


**Supplementary Figure 23**. hdWGCNA analysis on turtle gonads and network activity in rabbits. a) UMAP projection of the turtle gene network predicted by hdWGCNA. Genes are grouped based on their WGCNA module. The names for some top TFs per module are highlighted. b) Scatterplot displaying the kMEs of the genes belonging to the turtle *Pax2+* mesenchyme module in male rabbit vs male turtle. Highlighted are the androgen-synthesis genes, *PAX8*, *TBX1* and *PAX2*. In green if they are highly scored in the turtle, in blue if they are highly scored in the rabbit, in black if they are highly scored in both species and in grey if they do not surpass the threshold. c) Heatmap showing the ME scores of the turtle WGCNA modules in rabbit cell types. This implies that the genes belonging to this module are predominantly expressed in cell types with high MEs. d-e) Violin plot displaying the ME scores of the *Pax2*+ mesenchymal cells module in turtle cell types (d) and in rabbit cell types (e). Higher conservation in the rabbit SLCs is observed in comparison to the other cell types.

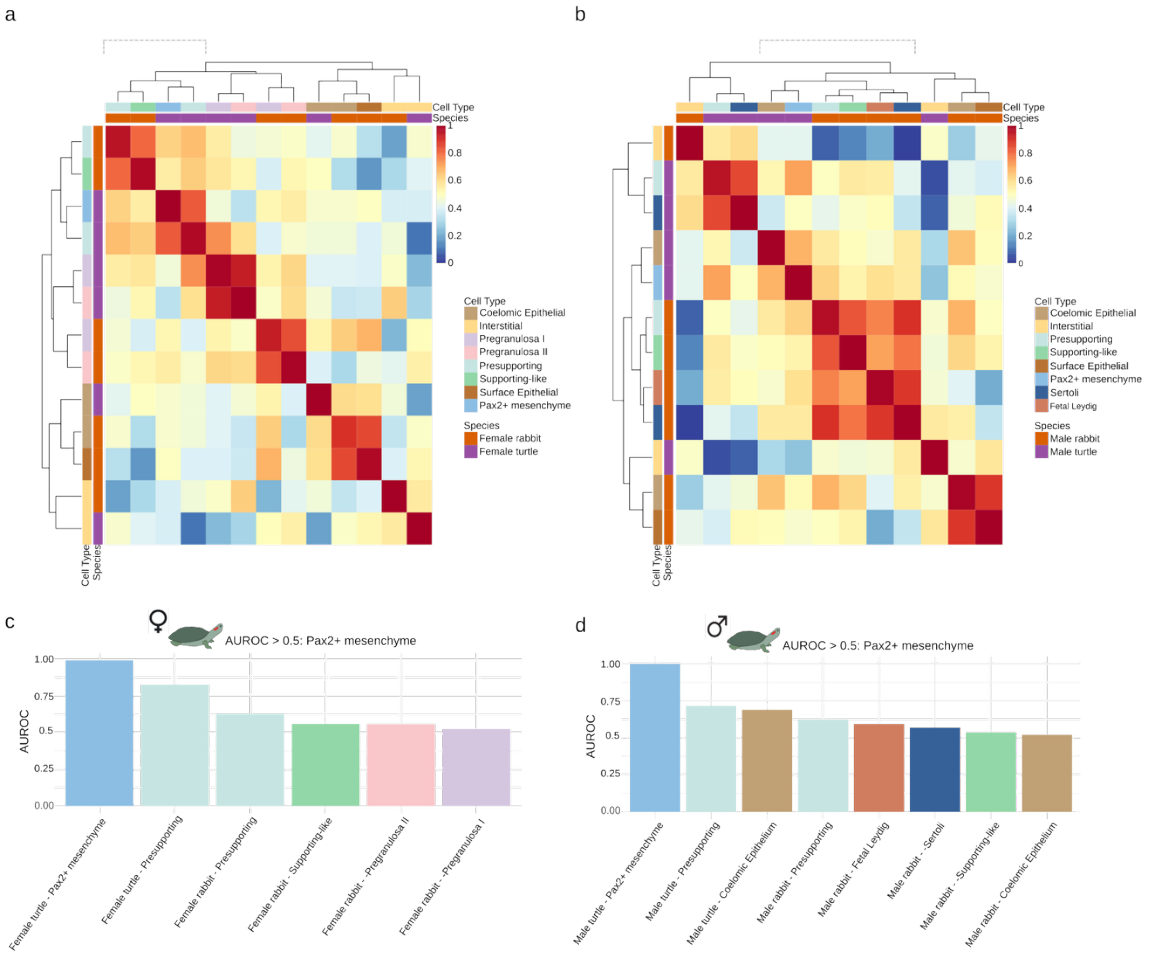


**Supplementary Figure 24**. Metaneighbor results. a-b) Heatmap displaying the AUROC scores from the MetaNeighborUS analysis in the female rabbit and female turtle cell types (a) and male rabbit and male turtle cell types (b). Highlighted with dashed lines the most similar cell types to rabbit SLCs. An AUROC value of 0.5 means that the transcriptional profile of the cell types are as similar as they would be by pure chance, therefore AUROC values higher than 0.5 indicate some degree of similarity between the cell types. c-d) Barplot highlighting the AUROC scores with respect to the turtle *Pax2*+ mesenchymal cells of female (c) and male (d) datasets.
